# Human brain glycoform co-regulation network and glycan modification alterations in Alzheimer’s disease

**DOI:** 10.1101/2023.11.13.566889

**Authors:** Qi Zhang, Cheng Ma, Lih-Shen Chin, Sheng Pan, Lian Li

## Abstract

Despite the importance of protein glycosylation to brain health, current knowledge of glycosylated proteoforms or glycoforms in human brain and their alterations in Alzheimer’s disease (AD) is limited. Here, we present a new paradigm of proteome-wide glycoform profiling study of human AD and control brains using intact glycopeptide-based quantitative glycoproteomics coupled with systems biology. Our study identified over 10,000 human brain N-glycoforms from nearly 1200 glycoproteins and uncovered disease signatures of altered glycoforms and glycan modifications, including reduced sialylation and N-glycan branching as well as elevated mannosylation and N-glycan truncation in AD. Network analyses revealed a higher-order organization of brain glycoproteome into networks of co-regulated glycoforms and glycans and discovered glycoform and glycan modules associated with AD clinical phenotype, amyloid-β accumulation, and tau pathology. Our findings provide novel insights and a rich resource of glycoform and glycan changes in AD and pave the way forward for developing glycosylation-based therapies and biomarkers for AD.

## Introduction

Alzheimer’s disease (AD) is the most common form of dementia, characterized pathologically by accumulation of amyloid-β (Aβ) plaques and tau neurofibrillary tangles in the brain (*1, 2*). Genome-wide association studies (GWAS) have identified over 40 genetic risk loci for AD (*3, 4*), and transcriptomic and proteomic analyses revealed RNA and protein expression changes that are much broader than Aβ and tau accumulation in AD brain (*3, 5*). However, DNA variations and RNA or protein level changes alone are insufficient to account for all aspects of AD pathobiology, as proteins are known to undergo posttranslational modifications that control their function (*6, 7*) and proteoforms generated from protein modifications add enormous diversity and complexity to the proteome (*7*). The most prevalent and complex form of protein modification is glycosylation, which produces vastly diverse arrays of glycosylated proteoforms or glycoforms (a glycoform is an isoform of a protein that differs only with respect to the number or type of attached glycan) to greatly expand the proteome and its functionalities (*8, 9*).

As a major type of glycosylation, N-linked glycosylation controls many biological processes through covalent attachment of glycans to asparagine residues of proteins (*9, 10*). Protein N-glycosylation involves a multi-step process that begins by oligosaccharyltransferase-catalyzed “en bloc” addition of a 14-sugar precursor Glc_3_Man_9_GlcNAc_2_ to asparagine residues within the sequon N-X-S|T (X ≠ P) of newly synthesized proteins in the lumen of endoplasmic reticulum (ER). The N-glycans are then processed and remodeled by coordinated actions of glycosyltransferases and glycosidases in the ER and Golgi compartments, producing a wide variety of glycoforms with distinct glycan compositions and structures, such as high-mannose-, hybrid-, and complex-type N-glycans (*9, 11*). The site-specific N-glycans on glycoproteins not only regulate protein folding, activity, trafficking, and subcellular localization but also serve as the sugar code to mediate bio-recognition and signal distinct cellular outcomes (*8, 12*).

N-glycan modifications of proteins are vital for development, maintenance, and function of the nervous system (*13, 14*). The importance of protein N-glycosylation to brain health is underscored by the findings that genetic mutations in glycosylation enzymes for N-glycan attachment or processing cause human congenital disorders with prominent neurological abnormalities (*15*). N-glycans on glycoproteins are increasingly recognized as functional effectors of genetic and epigenetic disease risk (*16*). Accumulating evidence points to a link between aberrant protein N-glycosylation and AD pathogenesis (*17–19*). Glycomic analyses of N-glycans detached from glycoproteins have revealed changes in global levels of N-glycans in AD brains (*20–22*), but the information of glycoproteins and glycosites bearing the N-glycans was lost. We recently performed an integrated proteomic and glycoproteomic study using ^18^O-tagging to label *in vivo* N-glycosylation sites and identified disease-associated N-glycoproteins and glycosites with altered N-glycosylation site occupancy in AD brains (*23*). While 1132 unique N-glycoproteins were analyzed, the study was limited by the use of a deglycosylation step in the workflow, resulting in the loss of information on glycans at each glycosite. A more recent intact glycopeptide-based glycoproteomic study reported qualitative analysis of site-specific N-glycan and glycoforms from 268 unique N-glycoproteins (defined by unique gene ID) in AD and control brains (*24*), but suffered from the low coverage of brain N-glycoproteome and the lack of quantitative information regarding the levels of site-specific N-glycans and glycoforms in human brain and their alterations in AD. Due to the inherent complexity of N-glycan modifications and technical challenges in intact glycoproteomics and associated data analysis, our current knowledge of system-wide changes in human brain glycoforms and site-specific N-glycan modifications in AD is very limited.

In this study, we established an integrated approach that combines intact glycopeptide-based quantitative glycoproteomics and systems biology for large-scale, in-depth analysis of glycoforms and site-specific N-glycan modifications in human brain and their changes in AD. Using this approach, we identified over 10,000 N-glycoforms from 1184 unique N-glycoproteins and obtained a system-level view of human brain glycoforms and their attached N-glycans. Our analyses reveal previously unknown changes in glycoforms and site-specific glycans in AD and uncover glycoform and glycan co-regulation networks in human brain and their alterations in AD. Our findings provide novel insights into the roles of N-glycan modifications in brain dysfunction in AD and establish a new framework of glycosylation-based networks and pathways for understanding and treating AD.

## Results

### Glycoproteomic landscape of AD and control brains revealed by intact glycoproteomics

To characterize human brain glycoproteome and its changes in AD, we established and optimized a mass spectrometry-based intact glycoproteomics platform for large-scale simultaneous profiling of glycoproteins, glycoforms, glycosites, and site-specific glycan compositions in brain samples. Our intact glycoproteomics workflow consisted of SDS-mediated protein extraction (*25*), filter-aided sample preparation (FASP) (*26*), glycopeptide enrichment by strong anion exchange and electrostatic repulsion-hydrophilic interaction chromatography (SAX-ERLIC) (*27*), liquid chromatography-tandem mass spectrometry (LC-MS/MS) with higher-energy collisional dissociation (HCD) using a stepped collision energy (SCE) setting (*28*), and MS data analysis using Byonic software (*29*) for identification of intact N-glycopeptides and their attached glycan compositions. We used this workflow to analyze the same sets of human dorsolateral prefrontal cortex tissue samples from 16 neuropathologically confirmed AD and control cases (table S1) as in our proteomics and glycosite-profiling studies (*23, 30*). In total, we identified 12,176 unique intact N-glycopeptides with attached N-glycans at the N-X-S|T (X ≠ P) sequon, corresponding to 10,731 unique N-glycoforms with 164 distinct N-glycan compositions at 2544 unique N-glycosites in 1184 unique N-glycoproteins (Fig. 1, A to D, and table S2). This dataset represents the largest glycoproteome dataset of human AD and control brains to date with site-specific N-glycan and glycoform information.

**Fig. 1.**
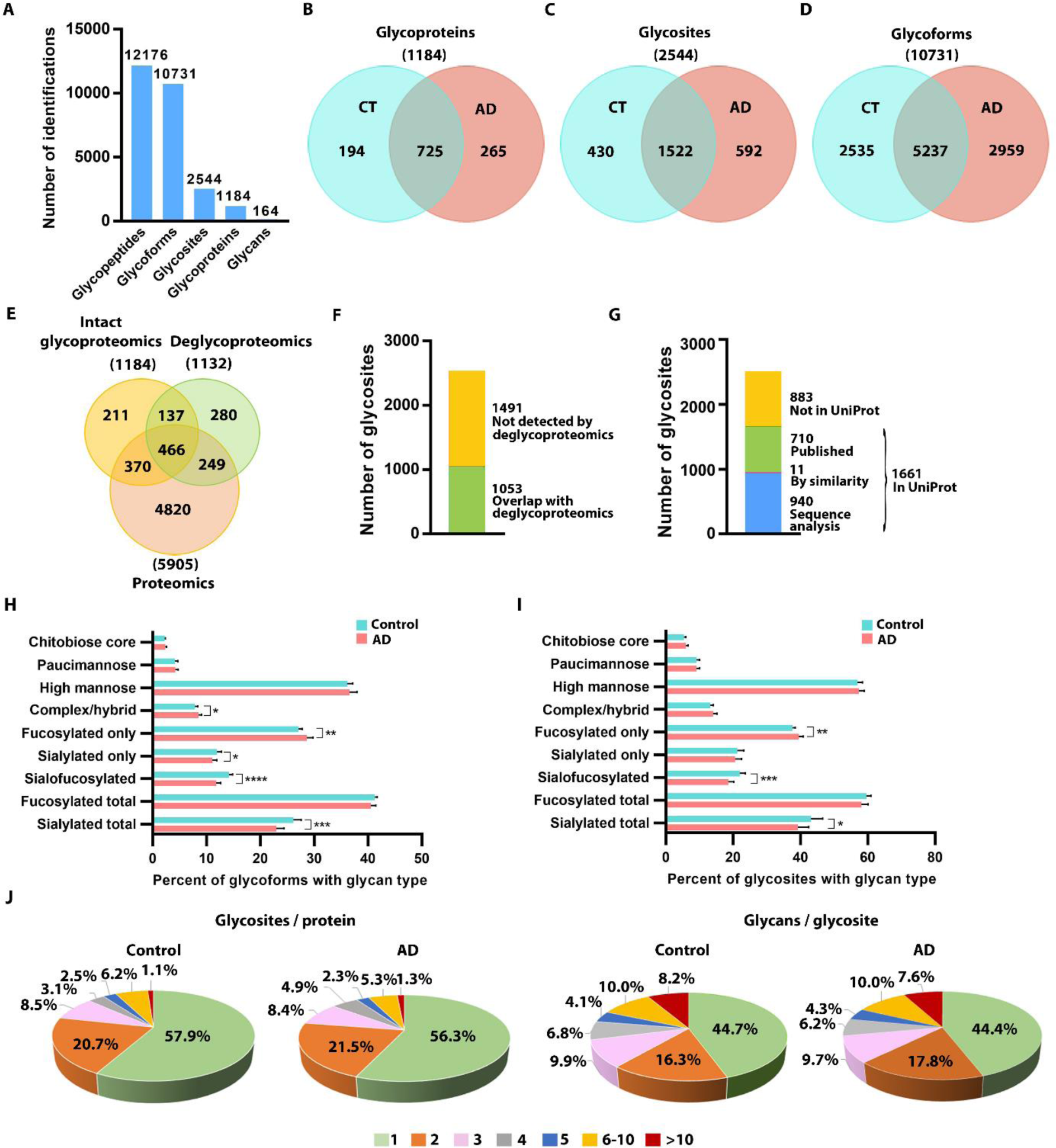
Intact glycopeptide-based glycoproteomics analysis of human AD and control brains. (**A**) The total number of unique N-glycopeptides, N-glycoforms, N-glycoproteins, N-glycosites, and N-glycan compositions identified in this study. (**B** to **D**) Comparisons of identified N-glycoproteins (B), N-glycosites (C), and N-glycoforms (D) in AD versus control (CT) brains. (**E**) Comparison of identified N-glycoproteins by intact glycoproteomics with proteins detected by proteomics and deglycoproteomics. (**F**) Overlap of identified N-glycosites with those detected by deglycoproteomics. (**G**) Match of the identified N-glycosites to Uniprot database of annotated N-glycosites. (**H** and **I**) Distribution of identified N-glycoforms (H) and N-glycosites (I) containing the indicated glycan type in AD and control brains. Data represent means ± SD (*n* = 8 cases per group). *, *P* < 0.05; **, *P* < 0.01; ***, *P* < 0.001; ****, *P* < 0.0001, unpaired two-tailed Student’s *t* test. (**J**) Pie charts showing the proportions of glycoproteins with the indicated number of *in vivo* N-glycosites per protein (macroheterogeneity) and glycosites with the indicated number of N-glycans per site (microheterogeneity) in AD and control brains.

Comparison of the intact glycoproteome dataset with our proteome profiling data from the same brain samples (*30*) showed that 348 glycoproteins (∼29%) of the 1184 glycoproteins were exclusively identified by intact glycoproteomics (Fig. 1E), demonstrating that our intact glycoproteomics workflow enables analysis of a subset of proteins that may otherwise remain undetected by global proteome profiling. As expected, the intact glycoproteome dataset was enriched for plasma membrane and extracellular proteins, lysosomal and ER lumenal proteins, neuronal and synaptic membrane proteins as well as receptors, ion channels, and cell adhesion molecules (fig. S1 and table S3). Our analyses (Fig. 1, E and F) showed that ∼51% of the N-glycoproteins and 41% of the N-glycosites identified in this study overlapped with those detected in our previous study (*23*) using a de-glycosylation-based glycoproteomics (deglycoproteomics) approach. The differences in the identifications of glycoproteins and glycosites between this work and our prior study are likely due to the differences in glycopeptide enrichment and MS fragmentation methods used in intact glycoproteomics versus deglycoproteomics analyses. Matching our intact glycoproteomic dataset to the UniProt database showed that 65% of the identified 2544 N-glycosites were annotated or predicted in the database (Fig. 1G). The current study provided experimental evidence for over 950 UniProt-predicted N-glycosites and identified over 880 new N-glycosites that have not been previously reported in UniProt (Fig. 1G). Comparison of our intact glycoproteomic data with the results of Suttapitugsakul et al (*24*) showed a substantial overlap in the identification of N-glycoproteins, N-glycosites, N-glycans, and N-glycoforms between these two studies (fig. S2). Notably, our intact glycoproteomics workflow achieved a greater coverage of brain glycoproteome, enabling analysis of additional 9597 N-glycoforms, 931 N-glycoproteins, 2080 N-glycosites, and 67 N-glycan compositions in this work (fig. S2).

Our glycoproteomic analysis identified 7772 N-glycoforms in control and 8196 N-glycoforms in AD, with 5237 N-glycoforms overlapping between control and AD brains (Fig. 1D, and table S2). We identified 164 N-glycan compositions in both control and AD brains (table S2) and categorized them into seven glycan types: high mannose, paucimannose, chitobiose core, undecorated complex/hybrid (neither fucosylated nor sialylated), fucosylated-only, sialylated-only, and sialofucosylated (sialylated and fucosylated) glycans. In control brains, ∼36% of the total N-glycoforms carried high-mannose-type glycans, ∼41% carried fucose-containing glycans (including both fucosylated-only glycans and sialofucosylated glycans), ∼26% carried sialic-acid-containing glycans (including both sialylated-only glycans and sialofucosylated glycans), ∼8% carried undecorated complex/hybrid (neither fucosylated nor sialylated) glycans, ∼4% carried paucimannose-type glycans, and ∼2% carried chitobiose-core-type glycans (Fig. 1H). We found that the percentages of sialylated-only glycoforms, sialofucosylated glycoforms, and total sialylated glycoforms were significantly decreased in AD brains, whereas the percentages of fucosylated-only glycoforms and undecorated complex/hybrid glycoforms were significantly increased in AD (Fig. 1H).

We identified 1952 N-glycosites in control and 2114 N-glycosites in AD brains, with 1522 N-glycosites shared by control and AD brains (Fig. 1C, and table S2). Approximately 57% of the total N-glycosites in control brains were observed with high-mannose glycans, ∼60% with fucose-containing glycans, 43% with sialic-acid-containing glycans, ∼13% with undecorated complex/ hybrid glycans, ∼9% with paucimannose glycans, and ∼5% with chitobiose-core-type glycans, (Fig. 1I). The percentages of sialofucosylated glycosites and total sialylated glycosites were significantly decreased in AD, whereas the percentage of fucosylated-only glycosites was significantly increased in AD (Fig. 1I). Our finding of disease-associated decreases in total sialylated glycosites (Fig. 1I) and sialylated glycoforms (Fig. 1H) indicates a reduction in overall sialylation in AD. For the 1522 shared N-glycosites, we analyzed the differences in the number of glycoforms at each glycosite between AD and controls for each of the major glycan types and performed unsupervised hierarchical clustering analysis to characterize site-specific N-glycosylation alterations in AD (fig. S3). The heatmap illustrated AD-associated changes in glycosylation patterns in ∼69% of the shared N-glycosites, whereas the remaining ∼31% of the glycosites showed no change in their glycosylation profiles in AD (fig. S3). For each glycan type, we observed increased glycosylation at some glycosites and decreased glycosylation at some other glycosites (fig. S3). Quantification of the net change in the number of glycoforms for each glycan type across the 1522 glycosites showed an overall decrease in sialylation in AD and an overall increase in modifications by high-mannose glycans, fucosylated-only glycans, and undecorated complex/hybrid glycans in AD (fig. S3).

Our analysis showed that more than half of the N-glycoproteins in control (∼58%) and AD (∼56%) brains were N-glycosylated at single glycosite, while about 10% were multiply glycosylated with five or more *in vivo* N-glycosylation sites (Fig. 1J). The average number of *in vivo* N-glycosites per glycoprotein was 2.1 in both control and AD brains, consistent with the result from our glycosite-profiling study (*23*). Our intact glycoproteomic analysis revealed extensive heterogeneity in the number of different N-glycans that modified each glycosite (Fig. 1J). Approximately 45% of the total glycosites were found with one glycan, whereas about 18% had more than five glycans and about 1% of the glycosites had more than 30 glycans in both control and AD brains (Fig. 1J). The identified glycosite with the highest degree of glycan heterogeneity was cell surface receptor CD47 N111 glycosite, which carried 78 glycans in control and 73 glycans in AD brains (table S2). The identified glycoprotein with the highest degree of glycoform diversity was neural cell adhesion protein contactin-1 (CNTN1), which had 199 glycoforms with 83 glycans across 8 glycosites in control brains and 194 glycoforms with 72 glycans across 8 glycosites in AD brains (table S2). On average, each glycosite had 4.0 glycans in control and 3.9 glycans in AD brains, and each glycoprotein had 8.5 glycoforms in control and 8.3 glycoforms in AD brains.

### Site-specific glycosylation profiles of AD-related proteins and risk factors

Our intact glycoproteomic analysis mapped site-specific N-glycans and N-glycoforms of a number of GWAS-identified AD risk factors (e.g., ABCA7, ACE, ADAM10, CLU, CNTNAP2, EPHA1, PLD3, SORL1, and TM2D3) and other AD-related proteins (e.g., tau, nicastrin, and LRP1) in human AD and control brains (Fig. 2, fig. S4, and table S2). The glycosylation profiles of these AD-relevant proteins demonstrated a broad range of glycan and glycoform diversity, with some proteins (e.g., ABCA7 and EPHA1) having only one glycoform carrying a single glycan at a single glycosite, while other proteins (e.g., CLU and LRP1) having over 80 glycoforms carrying 30 or more glycans across multiple glycosites (Fig. 2 and fig. S4). CLU (clusterin, also known as apolipoprotein J) is the third most significant risk factor for AD with an extracellular chaperone function (*31*). We identified a total of 88 CLU glycoforms carrying 60 different glycans across 4 glycosites in control brains and 104 CLU glycoforms carrying 67 glycans across 4 glycosites in AD brains, with a majority of glycoforms carrying fucosylated and/or sialylated glycans (Fig. 2 and table S2). Our analysis uncovered 26 AD-specific CLU glycoforms and 10 control-specific CLU glycoforms (Fig. 2 and table S2), revealing an involvement of CLU N-glycosylation aberrations in AD.

**Fig. 2.**
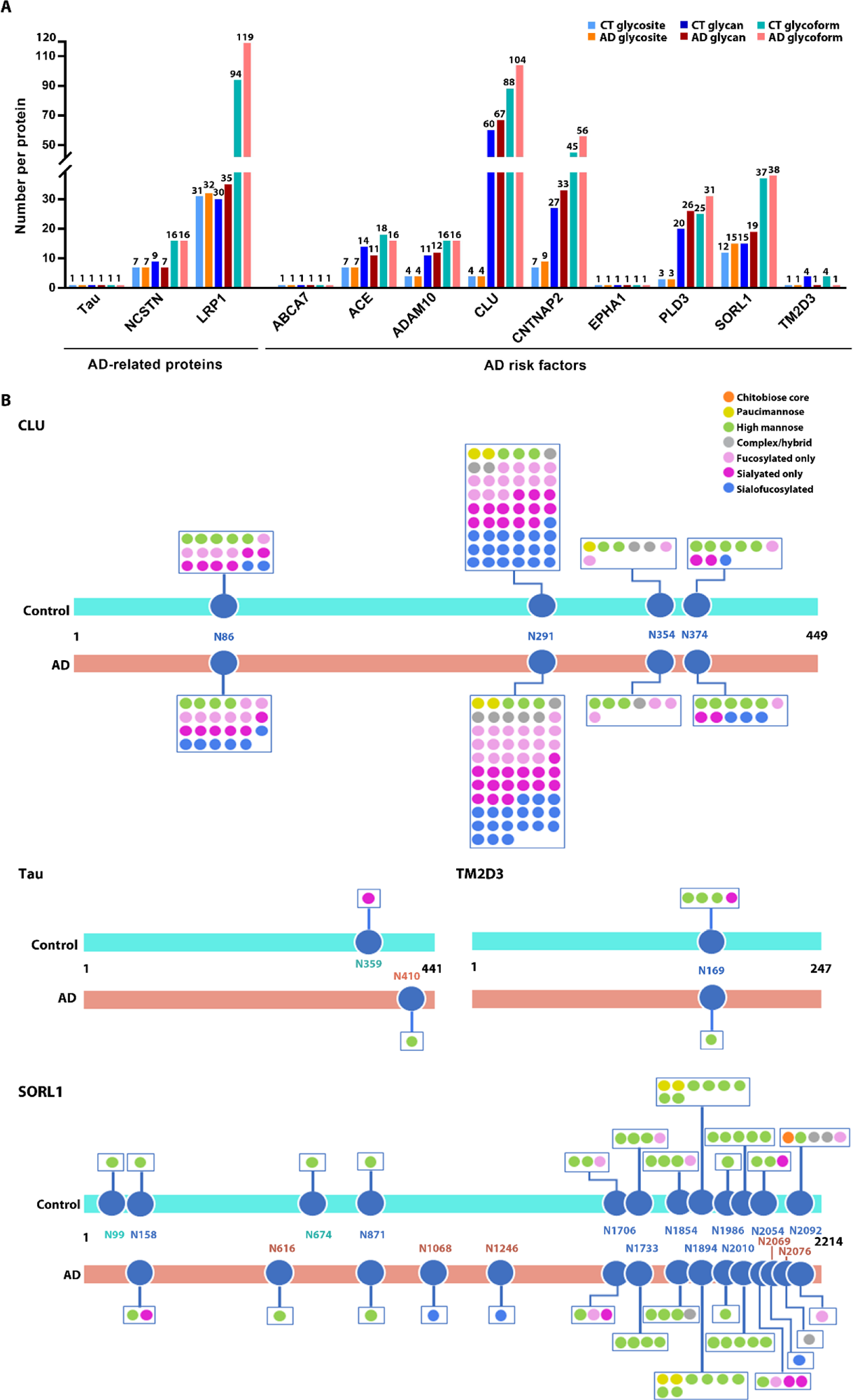
Site-specific glycan and glycoform mapping of AD-related proteins and risk factors. **(A)** Bar graphs showing the number of N-glycosites, N-glycan compositions, and N-glycoforms mapped to each of AD-related proteins and risk factors in AD and control (CT) brains. (**B**) Site-specific glycosylation patterns of CLU, tau, TM2D3, and SORL1 in AD compared to control brains. N-glycosites are indicated by the amino acid positions of glycosylated asparagine residues, with cyan- and pink-colored N-glycosites denoting the glycosites detected exclusively in control and AD brains, respectively.

SORL1 (sortilin-related receptor 1, also known as SORLA) is a cargo sorting receptor that has emerged as a major risk factor for both early-onset and late-onset AD (*1, 32*). Our analysis revealed 37 SORL1 glycoforms carrying 15 glycans across 12 glycosites in control brains and 38 SORL1 glycoforms carrying 19 glycans across 15 glycosites in AD brains, including 13 AD-specific glycoforms and 12 control-specific glycoforms, with a majority of glycoforms carrying high-mannose glycans (Fig. 2 and table S2). We identified five SORL1 glycosites (N616, N1068, N1246, N2069, and N2076) with a gain of N-glycosylation in AD and two SORL1 glycosites (N99 and N674) with a complete loss of N-glycosylation in AD (Fig. 2 and table S2). These findings, together with recent identification of SORL1 N674S mutation for a familial form of AD (*32, 33*), point to a pathogenic role of SORL1 N-glycosylation alterations in AD.

LRP1 (low-density lipoprotein receptor-related protein 1), a key regulator of Aβ and tau trafficking and AD neuropathology (*34, 35*), is the most heavily N-glycosylated protein that we identified in human brain (table S2). In total, 94 LRP1 glycoforms carrying 30 glycans across 31 glycosites were found in control brains and 119 LRP1 glycoforms carrying 35 glycans across 32 glycosites in AD brains, with a majority of glycoforms carrying high-mannose or fucosylated-only type glycans (fig. S4 and table S2). We identified 47 LRP1 glycoforms and 4 LRP1 glycosites exclusively in AD brains and 22 LRP1 glycoforms and 3 LRP1 glycosites only in controls (fig. S4 and table S2), highlighting an association of LRP1 N-glycosylation aberrations with AD.

We and others previously reported aberrant tau N-glycosylation in AD (*17–19, 23*), but the site-specific N-glycans and N-glycoforms of tau in AD remain unknown. Our analysis revealed that tau was modified at the N410 residue of tau441 isoform (corresponding to N727 of tau758 isoform) by a high-mannose-type glycan exclusively in AD but not controls (Fig. 2 and table S2), consistent with our prior finding of tau N410 as the N-glycosite in AD (*23*). We also detected a tau glycoform carrying a sialylated-only glycan at the N359 glycosite in control brains (Fig. 2 and table S2). In addition to CLU, SORL1, LRP1, and tau, our analysis revealed disease-associated changes in site-specific N-glycosylation patterns of several other AD-relevant proteins, including ACE, ADAM10, CNTNAP2, PLD3, TM2D3, and nicastrin (Fig. 2, fig. S4, and table S2). Together, these results provide strong support for the involvement of altered N-glycan modifications of the AD-related proteins and risk factors in AD pathophysiology.

### Reduced sialylation and N-glycan branching and elevated mannosylation and N-glycan truncation in AD

To further characterize protein N-glycan modifications and their alterations in AD, we performed quantitative analyses of intact glycopeptides based on their glycan type, monosaccharide composition, or other structural features such as branching to determine the relative abundance of glycan modification for each glycan category in individual brain samples from AD and control cases (Fig. 3, A to C, and table S4). We found that protein modification by high-mannose-type glycans is highly abundant in human brain, accounting for ∼49% of total abundance of all glycan modifications (Fig. 3A and table S4A). Five of the top seven most abundant N-glycans on brain proteins had compositions H5N2 to H9N2 corresponding to high-mannose glycan structures containing two core GlcNAc and five to nine mannose residues (Man5 to Man9 glycans), with H5N2 being the most abundant N-glycan at ∼21% of total glycan abundance in healthy human brains (Fig. 3C and table S4B). The other two of the top seven most abundant glycans had compositions H3N5F1 and H3N4F1 corresponding to bisecting GlcNAc-containing, core-fucosylated glycan structures (Fig. 3C and table S4B), consistent with the prior findings from glycomic analyses of N-glycans released from brain glycoproteins (*36, 37*). Our results revealed that the relative levels of H5N2, H3N5F1, and H3N4F1 modifications were significantly increased in AD, whereas H9N2 modification was significantly decreased in AD (Fig. 3C and table S4B).

**Fig. 3.**
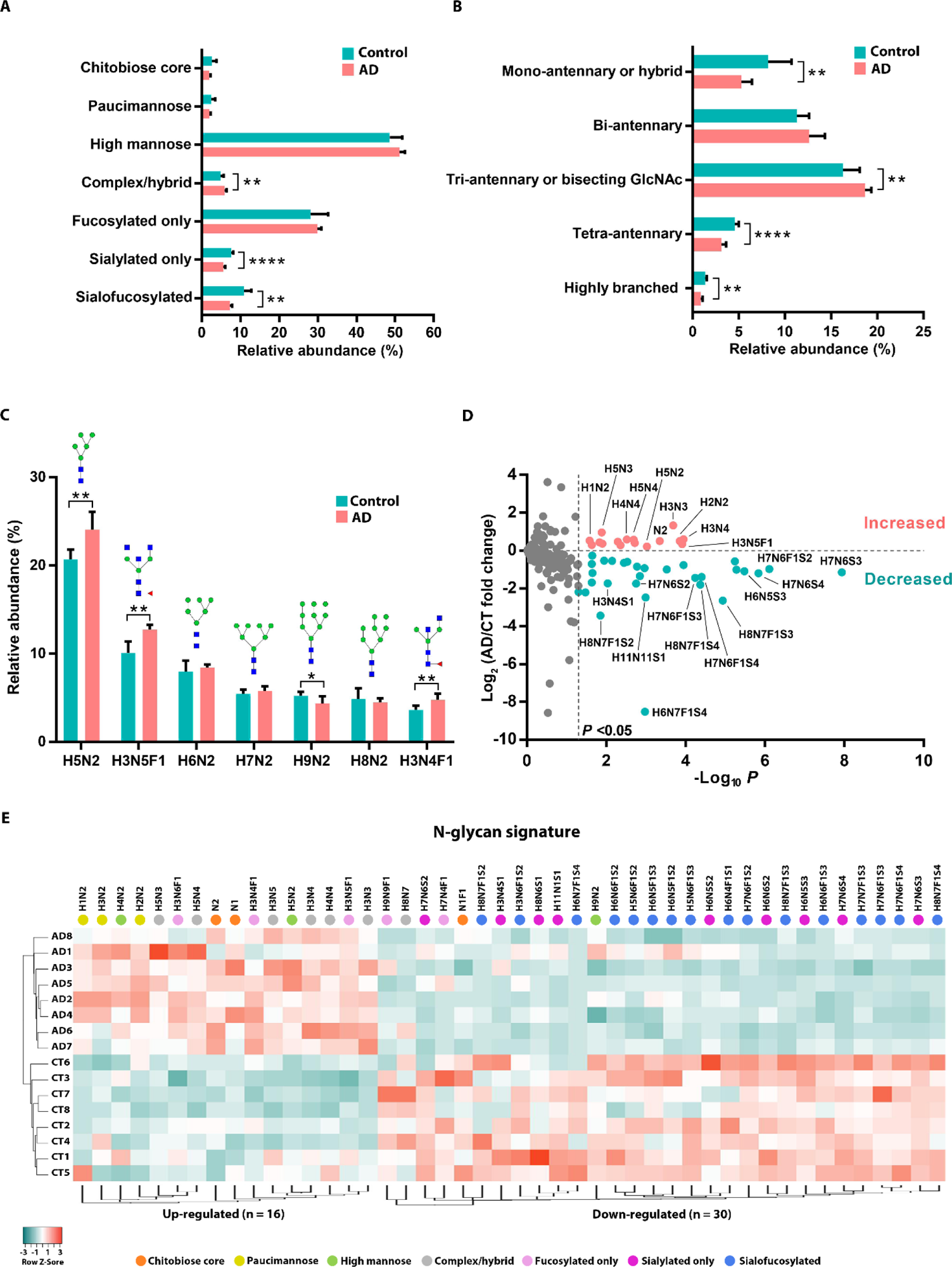
Disease-associated changes in overall levels of N-glycan modifications in AD. (**A to C**) Comparison of relative abundances of glycan modifications for the indicated glycan types (A), glycan branching categories (B), and top seven most abundant glycan compositions (C) in AD versus control brains. Corresponding glycan structures are shown for glycan compositions in (C). Data represent means ± SD (*n* = 8 cases per group). *, *P* < 0.05; **, *P* < 0.01; ***, *P* < 0.001; ****, *P* < 0.0001, unpaired two-tailed Student’s *t* test. (**D**) Volcano plot of fold changes in AD versus control (log_2_ FC) for 164 individual glycan compositions and corresponding –log_10_ *P* values showing identification of 46 significantly (*P* < 0.05) altered glycan modifications in AD with top 23 glycan modifications labeled. (**E**) Unsupervised hierarchical clustering based on glycan modification abundance profiles showing clear segregation of clinical cases into AD (top) and control (CT; bottom) groups and glycan modifications into up-regulated (left) and down-regulated (right) clusters in AD. Glycan composition key: H, hexose; N, N-acetylhexosamine; F, fucose; S, sialic acid.

Our quantitative analysis showed that in control brains, fucosylation (including modification by fucosylated-only and sialofucosylated glycans) and sialylation (including modification by sialylated-only and sialofucosylated glycans) occur at ∼39% and ∼19% relative abundance, respectively (Fig. 3A and table S4A). The sialylation in AD brains was reduced to ∼13% relative abundance, whereas the fucosylation level was not significantly changed in AD (Fig. 3A and table S4A). We also observed a significant increase in modification by complex/hybrid (neither fucosylated nor sialylated) glycans in AD (Fig. 3A and table S4A). Quantitative assessment of the extent of N-glycan branching revealed significant decreases in the overall levels of modifications by tetra-antennary glycans with six HexNAc residues and by highly branched and elongated glycans with more than six HexNAc residues in AD brains (Fig. 3B and table S4A), indicating decreased N-glycan branching in AD. In contrast, the relative level of modification by tri-antennary or bisecting GlcNAc glycans with five HexNAc residues was significantly increased in AD (Fig. 3B and table S4A), consistent with reported elevation in bisecting GlcNAc modification in AD (*38*).

We next performed differential abundance analysis of modifications by 164 individual glycan compositions and identified 46 glycan modifications that were significantly altered in AD compared to controls, including 30 glycan modifications with decreased abundance and 16 glycan modifications with increased abundance in AD (Fig. 3D and table S4, B and C). Unsupervised hierarchical clustering analysis based on glycan modification abundance profiles demonstrated that the identified 46 glycan modifications provide an N-glycan disease signature for distinguishing AD cases from the controls (Fig. 3E). The glycan clustering dendrogram showed a segregation of glycan modifications into down-regulated and up-regulated clusters in AD (Fig. 3E). We found that ∼83% of the identified down-regulated glycan modifications in AD were sialylation with a predominance of tetra-antennary or highly branched sialylated glycans (e.g., H8N6S1, H7N6F1S4, and H8N7F1S3), whereas none of up-regulated glycan modifications was sialylation (Fig. 3E and table S4C), providing additional evidence supporting an association of decreased sialylation and N-glycan branching with AD. The identified up-regulated glycan modifications in AD (Fig. 3E and table S4C) include mannosylation by H4N2 and H5N2 glycans, modification by undecorated complex/hybrid glycans with relatively simple, neutral structures (e.g., H3N3, H5N3, H4N4, and H5N4), and modification by bisecting GlcNAc-containing core-fucosylated glycans (e.g., H3N4F1, H3N5F1, and H3N6F1). We also observed increased modifications by chitobiose-core-type glycans N1 and N2 and by paucimannose glycans H1N2, H2N2, and H3N2 (Fig. 3E and table S4C), indicating elevated N-glycan truncation in AD.

### AD-associated protein glycoforms with altered site-specific N-glycan modifications

Next, we assessed disease-associated changes at the levels of individual glycoforms by performing differential glycoform abundance analysis (Fig. 4A and table S5A) and identified 556 glycoforms with significantly altered levels (>1.3-fold change, *P* < 0.05) in AD compared to control brains, including 421 glycoforms from 149 glycoproteins with increased glycoform abundance and 135 glycoforms from 58 glycoproteins with decreased glycoform abundance in AD (Fig. 4, A and B, and table S5B). Comparison of glycoform abundance changes with the corresponding protein abundance changes measured from the same brain samples (*30*) showed that most of the observed changes in glycoform abundance in AD were caused by changes in site-specific N-glycan modification levels but not by altered protein expression levels (Fig. 4C, fig. S5, and table S5B). We found that 449 glycoforms had >1.3-fold changes in site-specific N-glycan modification levels in AD versus controls, including 337 glycoforms from 111 glycoproteins with increased N-glycan modification levels and 112 glycoforms from 52 glycoproteins with decreased N-glycan modification levels in AD (Fig. 4C and table S5B). Unsupervised hierarchical clustering analysis showed that the identified 449 glycoforms segregated into down-regulated and up-regulated clusters and they provide a glycoform signature for distinguishing AD cases from the controls (Fig. 4D). We observed an enrichment of sialylated-only and sialofucosylated glycans in the down-regulated cluster and an enrichment of fucosylated-only, paucimannose-type, and high-mannose-type glycans in the up-regulated cluster (Fig. 4D and table S5B). The identified N-glycans attached to disease-associated glycoforms (Fig. 4D and table S5B) provide a site-specific glycan signature for AD.

**Fig. 4.**
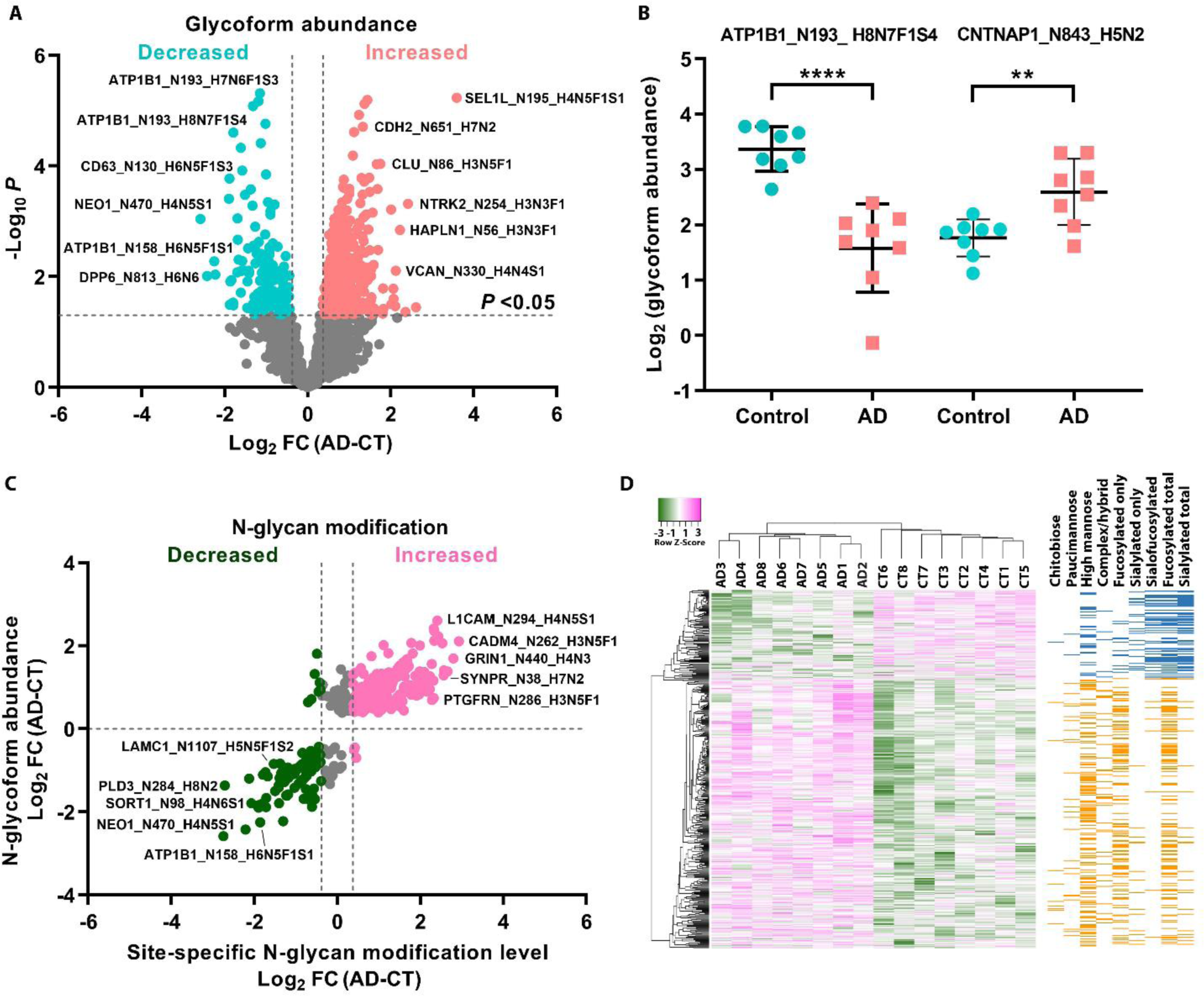
Identification of AD-associated protein glycoforms with altered site-specific N-glycan modifications. (**A**) Volcano plot of individual glycoform abundance fold changes in AD versus control (log_2_ FC) and corresponding–log_10_ *P* values showing identification of 556 significantly altered (> 1.3-fold change, *P* < 0.05) glycoforms in AD with top 12 glycoforms labeled. (**B**) Examples of identified glycoforms with decreased or increased abundance in AD. Data are shown as means ± SD (*n* = 8 cases per group). **, *P* < 0.01; ****, *P* < 0.0001, unpaired two-tailed Student’s *t* test. (**C**) Scatter plot showing at least ±1.3-fold changes in site-specific glycan modifications for the majority of the identified AD-associated glycoforms with top 10 glycoforms labeled. (**D**) Unsupervised hierarchical clustering of individual clinical cases based on the identified 449 glycoforms with altered site-specific glycan modifications. Glycoforms are displayed as rows with corresponding glycan types shown in the *left* heatmap.

Beside the glycoforms with altered glycan modification levels, our analyses also identified 179 glycoforms from 105 glycoproteins with a gain of a site-specific N-glycan modification in AD and 49 glycoforms from 32 glycoproteins with a loss of a site-specific N-glycan modification in AD (Fig. 5, A to C, and table S5C), providing additional site-specific glycoform and glycan signatures for AD. In total, we identified 516 hyperglycosylated glycoforms from 178 hyperglycosylated proteins with increased or a gain of site-specific N-glycan modification in AD and 161 hypoglycosylated glycoforms from 71 hypoglycosylated proteins with decreased or a loss of site-specific N-glycan modification in AD (Fig. 5, A to C, and table S6, A and B). We found that both hyperglycosylation and hypoglycosylation datasets contain glycoforms and glycoproteins with altered paucimannosylation, oligomannosylation, fucosylation, and/or sialylation (Fig. 5, A to C, fig. S6, and table S6).

**Fig. 5.**
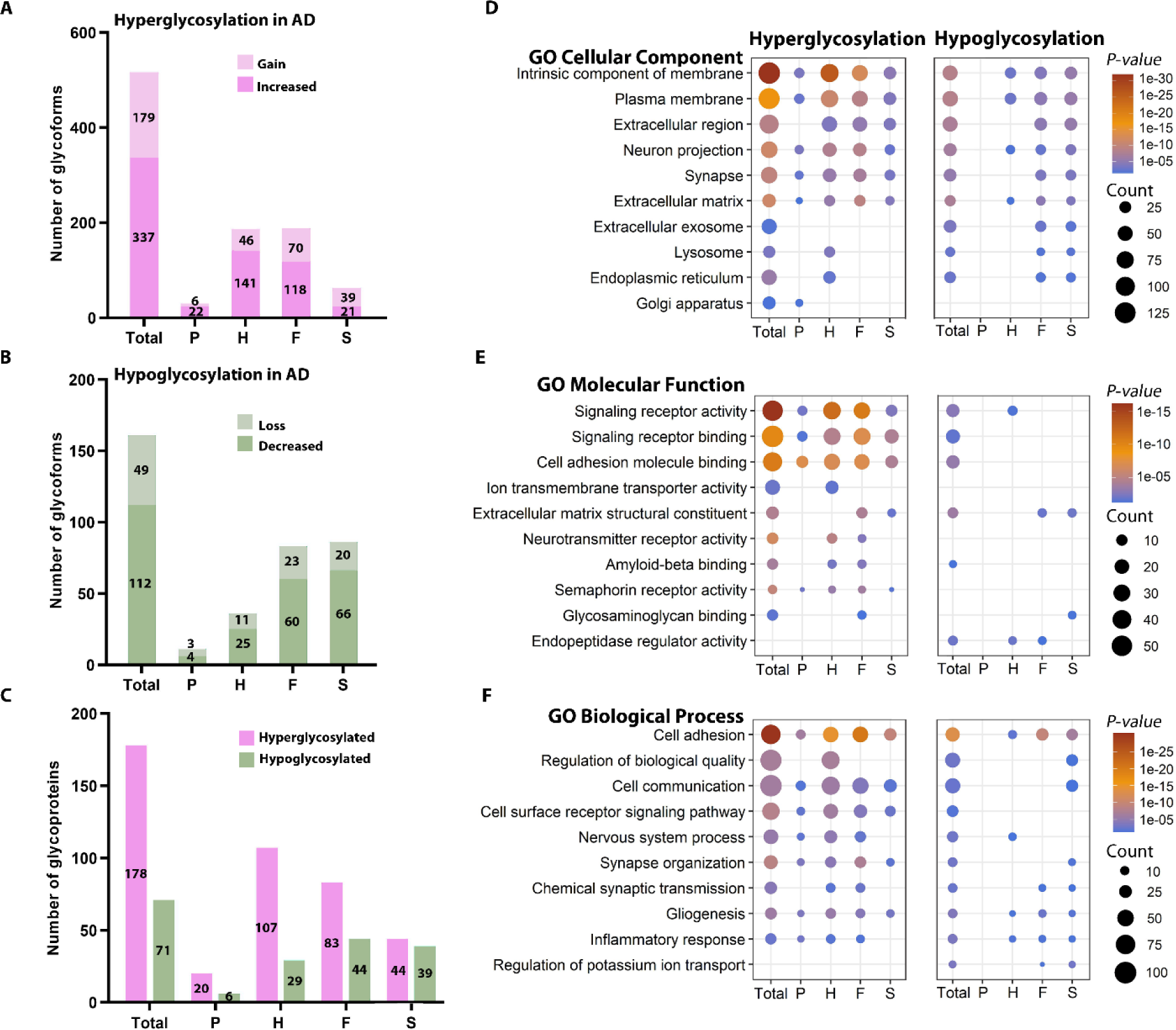
Analysis and annotation of glycoforms and glycoproteins with altered site-specific N-glycan modifications in AD. (**A** to **C**) Hyperglycosylation includes glycoforms (A) and glycoproteins (C) with increased site-specific N-glycan modification and/or a gain of site-specific N-glycan modification in AD, whereas hypoglycosylation includes glycoforms (B) and glycoproteins (C) with decreased site-specific N-glycan modification and/or a complete loss of site-specific N-glycan modification in AD. Numbers indicate the total number or the number of hyperglycosylated or hypoglycosylated glycoforms or glycoproteins carrying the following glycan types: P, paucimannose; H, high mannose; F, fucose-containing glycan; and S, sialic-acid-containing glycan. **(D** to **F)** GO cellular component, molecular function, and biological process categories enriched in AD hyperglycosylation or hypoglycosylation dataset are shown with Benjamini-Hochberg FDR-corrected *P* values. Count indicates the number of glycoproteins per GO term.

We then performed gene ontology (GO) enrichment analyses of the hyperglycosylation and hypoglycosylation datasets to gain insights into the cellular functions and processes that were affected by dysregulated N-glycan modifications in AD (Fig. 5, D to F, and table S7, A and B). We found that both hypoglycosylation and hyperglycosylation datasets were significantly enriched for plasma membrane and extracellular proteins involved in cell adhesion, cell surface receptor signaling, nervous system process, and extracellular matrix function. However, only AD-associated hyperglycosylation, but not hypoglycosylation, was significantly linked to GO terms of neurotransmitter receptor activity and semaphorin receptor activity (Fig. 5E and table S7, A and B), whereas hypoglycosylation in AD, but not hyperglycosylation, was enriched for GO categories of endopeptidase regulator activity and regulation of potassium ion transport (Fig. 5, E and F). Our analyses revealed multiple cellular functions that were differentially impacted by different types of altered glycan modifications in AD (Fig. 5, D to F, and table S7, C to J). For example, hyperoligomannosylation, hypofucosylation, and hyposialylation in AD, but not other types of glycan modification changes, were significantly associated with ER and lysosome, whereas hyperpaucimannosylation was selectively associated with Golgi apparatus (Fig. 5D). Hyperoligomannosylation and hyperfucosylation in AD were found to specifically associate with amyloid-beta binding (Fig. 5E), and hyperoligomannosylation and hyposialylation in AD were preferentially linked to regulation of biological quality (Fig. 5F).

### Glycan modification co-regulation network analysis uncovers glycan modules linked to AD pathology

To gain systems-level insights into glycan modifications in human brain and their changes in AD, we performed glycan modification co-regulation network analysis by using weighted correlation network analysis (WGCNA) algorithm (*39*) to construct a glycan network with nodes representing glycan modifications connected with edges defined by the connectivity based on pairwise correlation patterns of glycan modification abundance profiles of 164 glycan compositions (Fig. 6A). By applying the scale-free topology criterion (*39*), we generated a human brain glycan network consisting of 8 modules of strongly co-regulated glycan modifications (Fig. 6, A and B, and table S8). These glycan modules (GNMs), color coded using the WGCNA convention (*39, 40*), were labeled according to module size from GNM1 (the largest module with 52 glycans) to GNM8 (the smallest module with 5 glycans) (Fig. 6B and table S8). We computed an eigenglycan for each module as the module representative and performed module-trait association analyses to assess the correlation relationships between each module eigenglycan and AD-relevant phenotypic traits and other characteristics of brain samples (fig. S7). We identified one positively correlated glycan module (GNM2) and two negatively correlated glycan modules (GNM1 and GNM4) with significant association to AD status, Aβ pathology (CERAD score), and/or neurofibrillary tangle pathology (Braak stage), but no association to age, gender, ApoE genotype, or postmortem interval (Fig. 6B and fig. S7). We found that AD-positively correlated module GNM2 had significantly increased module glycan modification levels in AD, whereas AD-negatively correlated modules GNM1 and GNM4 had significantly decreased module glycan modification levels in AD (Fig. 6C).

**Fig. 6.**
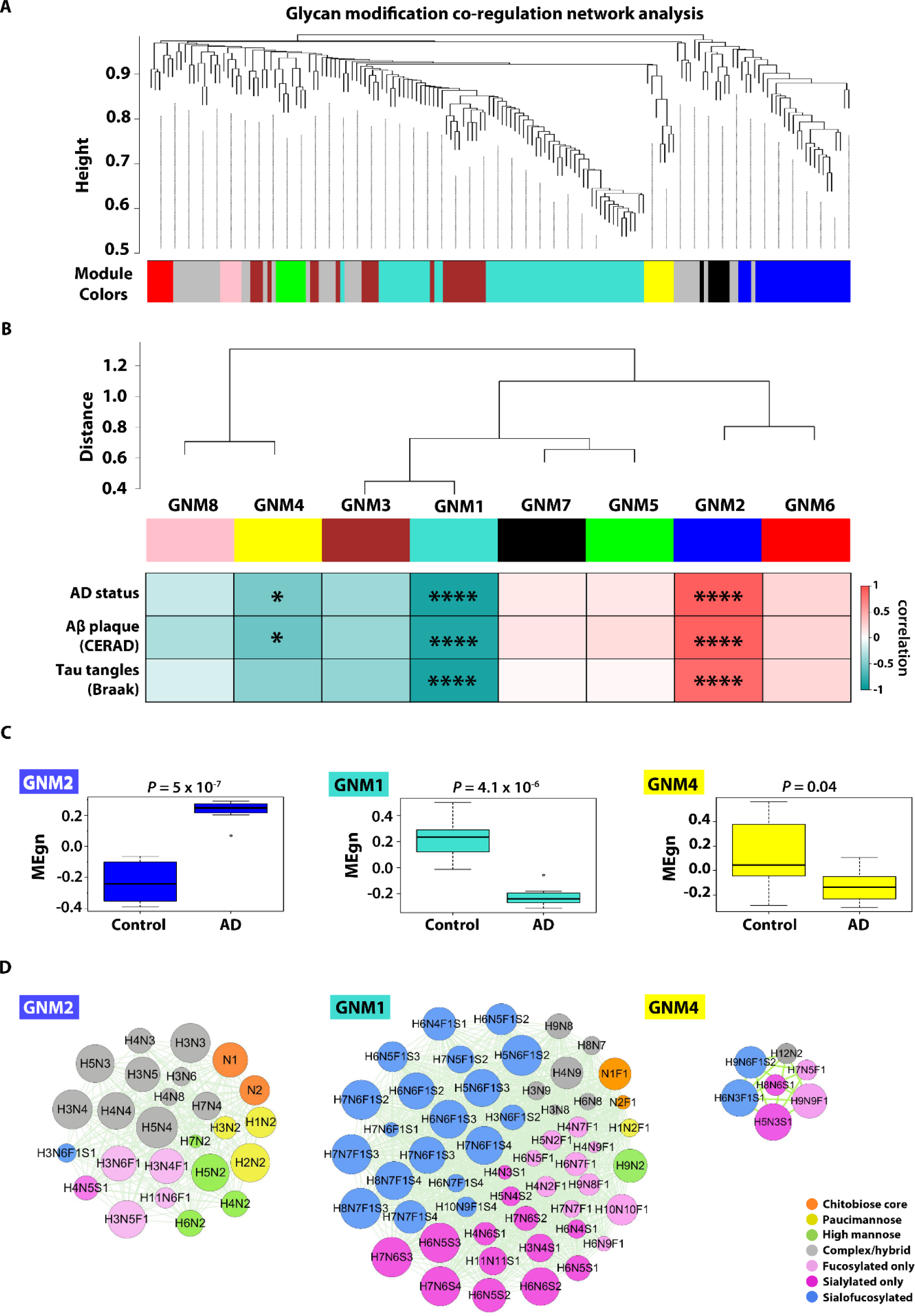
Glycan modification co-regulation network analysis and identification of AD-related glycan modules. (**A**) Glycan WGCNA cluster dendrogram with dynamic tree cut revealing 8 modules (color-coded) of co-regulated N-glycan modifications. (**B**) Inter-modular connectivity of the identified 8 glycan modules. Correlations of each glycan module with AD status, Aβ pathology (CERAD score), or neurofibrillary tangle pathology (Braak stage) are shown by Biweight midcorrelation coefficients (bicor *r*-values) in the heatmap, with significant positive (red) or negative (green) correlations marked by asterisks. *, *P* < 0.05; ****, *P* < 0.0001. (**C**) Box plots of module eigenglycan (MEgn) values in AD and control cases for each AD-related module, with differences in MEgn values between AD and control shown by Kruskal–Wallis test *P* values. (**D**) Glycan co-regulation network plots of AD-associated modules GNM2, GNM1, and GNM4, with node representing individual glycan composition, node color for glycan modification type, and node size in proportion to node connectivity. Glycan composition key: H, hexose; N, N-acetylhexosamine; F, fucose; S, sialic acid.

Our analyses showed that the GNM2 module, which has a strong positive correlation to AD neuropathology (CERAD *r* = 0.91, *P* = 1 x 10^-6^; Braak *r* = 0.85, *P* = 4 x 10^-5^), contained 25 glycan modifications (Fig. 6D), including modifications by oligomannose glycans (H4N2, H5N2, H6N2, and H7N2), undecorated complex/hybrid glycans with relatively simple, neutral structures (e.g., H3N3, H3N4, and H5N4), bisecting GlcNAc-containing core-fucosylated glycans (e.g., H3N4F1, H3N5F1, and H3N6F1), paucimannose glycans (H1N2, H2N2, and H3N2), and chitobiose-core glycans (N1 and N2). In contrast, the GNM1 module with a strong negative correlation to AD neuropathology (CERAD *r* = -0.87, *P* = 1 x 10^-5^; Braak *r* = -0.9, *P* = 2 x 10^-6^) contained 52 glycan modifications (Fig. 6D), which primarily consisted of modifications by sialofucosylated glycans (e.g., H5N6F1S2, H7N7F1S3, and H8N7F1S3), sialylated-only glycans (e.g., H6N5S3, H7N6S3, and H7N6S4), and highly branched glycans (e.g., H4N9, H10N10F1, and H11N11S1). Interestingly, we found co-segregation of core-fucosylated glycans N1F1, N2F1, H1N2F1, H4N2F1, and H5N2F1 in AD-negatively correlated GNM1 module, whereas their non-fucosylated counterparts (N1, N2, H1N2, H4N2, and H5N2) were co-segregated in AD-positively correlated GNM2 module (Fig. 6D), indicating an association of reduced core-fucosylation of these glycans with AD. Our analyses also showed that the GNM4 module with negative correlation to Aβ pathology but no correlation to tau tangle pathology (Fig. 6D) contained 7 glycan modifications, including modifications by sialofucosylated glycans H6N3F1S1 and H9N6F1S2, sialylated-only glycans H5N3S1 and H8N6S1, and highly branched glycan H9N9F1, suggesting an Aβ-specific negative effect on these glycan modifications. The predominant presence of sialylated glycans (including sialylated-only and sialofucosylated glycans) and highly branched N-glycans in AD-negatively correlated GNM1 and GNM4 modules further supports the involvement of decreased sialylation and N-glycan branching in AD pathophysiology.

### Protein glycoform co-regulation network analysis reveals AD-associated glycoform modules

To elucidate the relationships among protein glycoforms with different site-specific glycan modifications in AD, we performed glycoform co-regulation network analysis using WGCNA to construct a glycoform network with nodes representing glycoforms connected with edges defined by the connectivity based on pairwise correlation patterns of glycoform abundance profiles (Fig. 7A). Our network analysis revealed an organization of human brain glycoproteome into a network of 21 modules of co-regulated glycoforms (Fig. 7, A and B, and table S9). These glycoform modules (GFMs) were labeled from GFM1 (the largest module with 440 glycoforms from 159 glycoproteins) to GFM21 (the smallest module with 36 glycoforms from 16 glycoproteins) (Fig. 7B and table S9). In the glycoform network, different glycoforms from a single protein were often found in different modules, such as the differential distribution of GABA-B receptor subunit GABBR1 glycoforms to GFM2 (GABBR1_N440_H5N2 and GABBR1_N440_H3N5F1 glycoforms), GFM4 (GABBR1_N482_H5N2 glycoform), GFM8 (GABBR1_N440_H4N4F1 glycoform), GFM14 (GABBR1_N502_H4N5S1 glycoform), and GFM21 (GABBR1_N440_ H5N4F1 glycoform) modules (table S9). The colocalization of some glycoforms in the same module (e.g., the presence of two GABBR1 glycoforms in the GFM2 module) suggests coordinated regulation of these glycoforms and their site-specific glycan modifications, whereas the partition of glycoforms into separate modules (e.g., the distribution of GABBR1 glycoforms in the GFM2, GFM4, GFM8, GFM14 and GFM21 modules) may reflect differential regulation of site-specific glycan modifications or different cellular or subcellular distributions of distinct glycoforms.

**Fig. 7.**
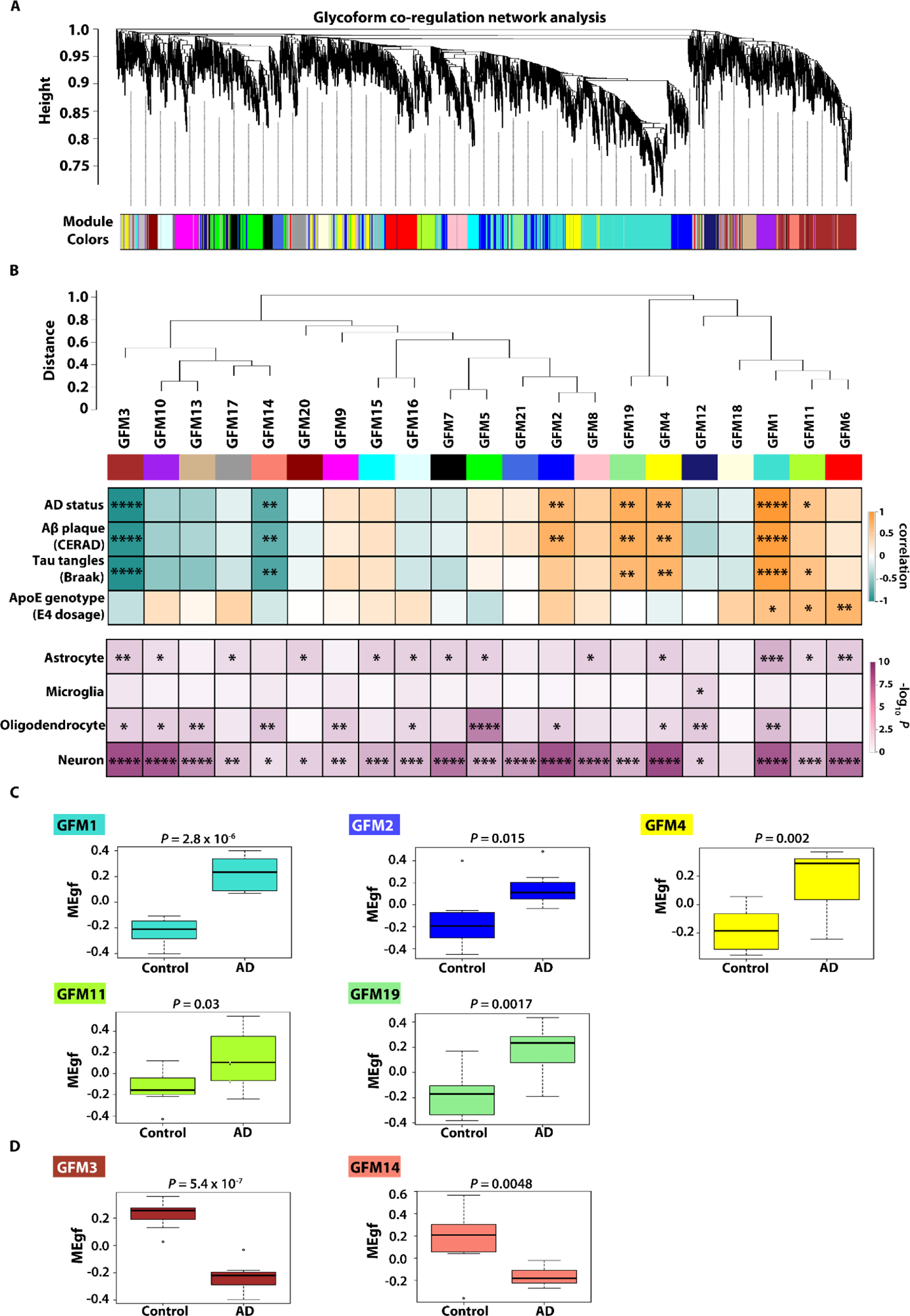
Protein glycoform co-regulation network analysis and identification of AD-associated glycoform modules. (**A**) Glycoform WGCNA analysis identifies 21 modules of co-regulated protein glycoforms. (**B**) Inter-modular connectivity of 21 glycoform co-regulation modules. Heatmap at the *top* shows the correlations (bicor *r*-values) of each glycoform module with the indicated AD-relevant traits, with significant positive (orange) or negative (green) correlations marked by asterisks. Heatmap at the *bottom* shows the enrichment for markers of astrocyte, microglia, oligodendrocyte, or neuron in each module with FDR-adjusted *P* values. *, *P* < 0.05; **, *P* < 0.01; ***, *P* < 0.001; ****, *P* < 0.0001. (**C**) Box plots of module eigenglycoform (MEgf) values in AD and control cases for each AD-associated module, with differences in MEgf values between AD and control shown by Kruskal–Wallis test *P* values.

We performed module-trait association analyses using module eigenglycoforms as the module representatives and identified seven glycoform modules with significant correlations to AD status, Aβ CERAD score, and/or tau Braak stage, including five positively correlated modules (GFM1, GFM2, GFM4, GFM11, and GFM19) and two negatively correlated modules (GFM3 and GFM14) (Fig. 7B and fig. S8). None of the seven glycoform modules showed significant correlation to age, sex, or postmortem interval (fig. S8), indicating that the identified AD-association of these modules was not due to the potential confounding factors. We found that GFM1 and GFM11 modules also showed positive correlations with the ApoE4 allele dosage (Fig. 7B and fig. S8), in line with the dosage-dependent role of ApoE4 allele for increasing AD risk (*41*). In addition, we identified an ApoE4-positively correlated module (GFM6) with no association to AD phenotypes (fig. S8), indicating that some of ApoE4-induced changes in site-specific glycan modifications are unrelated to AD pathology. Our analysis showed that the module glycoform abundance profiles for all five AD-positively correlated modules were significantly increased in AD, whereas those for the two AD-negatively correlated modules were significantly decreased in AD (Fig. 7C).

We then assessed the enrichment of cell-type markers for different brain cells in glycoform modules and found that six of the seven AD-associated glycoform modules (GFM1, GFM2, GFM3, GFM4, GFM11, and GFM19) were highly enriched with neuronal cell markers, although some of these modules were also enriched with astrocyte and/or oligodendrocyte markers (Fig. 7B). By contrast, AD-associated GFM14 module was preferentially enriched with oligodendrocyte markers (Fig. 7B). Analysis of inter-modular relationships using pairwise correlations between module eigenglycoforms revealed a higher-order organization of closely interconnected glycoform modulesinto superclusters (Fig. 7B). The colocalization of AD-negatively correlated GFM3 and GFM14 modules in a supercluster (Fig. 7B) suggests close relationships among the glyco-pathways and/or biological processes associated with these two negatively correlated glycoform modules. We found that four of the five AD-positively correlated modules (GFM1, GFM4, GFM11, and GFM19) were co-localized in another supercluster (Fig. 7B), indicating that the corresponding glyco-pathways and/or biological processes for these positively correlated modules may also be functionally related and/or coordinately regulated in AD.

Topological structures and membership information of the seven AD-associated glycoform modules are provided in Fig. 8A and table S9. Given the central role of hub nodes in determining a network module’s architecture and function (*42, 43*), we identified hub glycoforms for each module using the intramodular connectivity and depicted top 10 most-connected hub glycoforms in the center of each network plot (Fig. 8A). We assessed the glycan characteristics of each AD-related module by analyzing the distributions of site-specific glycan types and glycan compositions attached to glycoforms in each module (Fig. 8, B and C) and found that each AD-related module had a distinctive site-specific glycan modification pattern, which reflects module-specific co-regulation among glycoforms and their site-specific glycan modifications in the module. For example, GFM1 module was enriched for glycoforms carrying chitobiose-core, paucimannose, oligomannose, or fucosylated-only glycans and underrepresented for glycoforms with sialylated-only or sialofucosylated glycans, whereas GFM3 module was enriched for sialofucosylated and sialylated-only glycoforms but underrepresented for glycoforms bearing other glycan types (Fig. 8B). Our analyses revealed that enrichment of oligomannosylated glycoforms and depletion of sialofucosylated glycoforms is a common feature shared by all five AD-positively correlated modules, and the opposite was true for AD-negatively correlated modules, which were enriched for sialofucosylated glycoforms and depleted for oligomannosylated glycoforms (Fig. 8B).

**Fig. 8.**
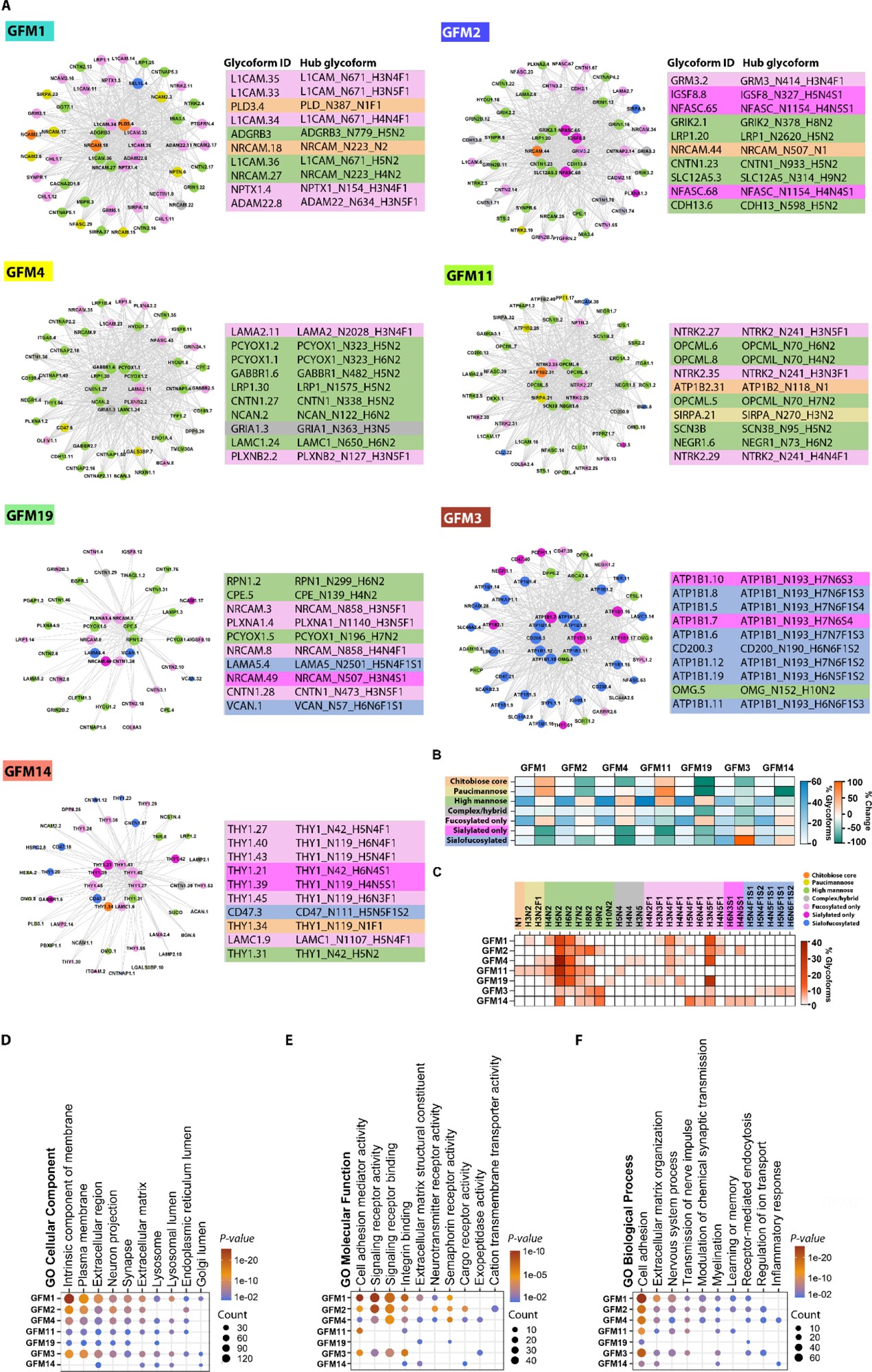
Biochemical and functional characteristics of AD-associated glycoform module networks. (**A**) Glycoform co-regulation network plots of AD-associated modules, with node representing individual glycoform, node color for site-specific glycan modification type, and node size in proportion to kME-based intramodular connectivity. A maximum of 50 glycoforms are shown for each module, with top 10 hub glycoforms in the center of each network plot. The glycoform ID and full name (protein_N-glycosite_glycan composition) for top 10 hub glycoforms are indicated next to the module network plot, with glycoforms color-coded for site-specific glycan types. The full list of glycoforms in each module and their kME values are provided in table S9. (**B**) Heatmap shows the percentages of glycoforms with the indicated glycan types in each AD-associated module (% Glycoform scale bar) and the net change in the percentage of glycoforms carrying each glycan type in each module versus the percentage of glycoforms carrying the same glycan type in the control brain (% Change scale bar). (**C**) Heatmap shows the percentages of glycoforms carrying each of top 10 most frequent glycan compositions in each AD-associated module. **(D** to **F)** GO cellular component, molecular function, and biological process categories enriched in each AD-associated module are shown with Benjamini-Hochberg FDR-corrected *P* values. Count indicates the number of glycoproteins per GO term.

### Glycoform module analyses uncover multiple CNS processes altered by dysregulated glycan modifications in AD

We performed GO enrichment analyses and found that the identified AD-related glycoform modules had overlapping yet distinct biochemical and functional characteristics (Fig. 8, D to F, and table S10). All seven disease-related modules were prominently associated with cell adhesion process with an overrepresentation of glycoforms from cell adhesion proteins, including neural cell adhesion molecules (e.g., NCAM1, NCAM2, NRCAM, L1CAM, and CHL1/L1CAM2), synaptic cell adhesion molecules (CADM1/SynCAM and CADM2/SynCAM2), IgLON family of cell adhesion molecules (OPCML, LSAMP, and NEGR1), adhesion G protein-coupled receptors (ADGRB1/BAI1, ADGRB3/BAI3, and ADGRL1/ latrophilin-1), neurexin-1 (NRXN1), teneurin-4 (TENM4), neuronal pentraxin-1 (NPTX1), neuronal pentraxin receptor (NPTXR), contactins (CNTN1, CNTN2, CNTN3, and CNTN4), contactin-associated proteins (CNTNAP1/Caspr1, CNTNAP2/Caspr2, CNTNAP4/Caspr4, and CNTNAP5/Caspr5), cadherins (CDH2, CDH10, and CDH13), protocadherins (PCDH1, PCDH9, and PCDH17), neurofascin (NFASC), neogenin (NEO1), neuroplastin (NPTN), ADAM22, LGI1, CD47, CD200, SIRPA, JAM3, and/or THY1 (Fig. 8, E and F, and table S9). The identification of site-specific glycoforms from cell adhesion proteins as top hub glycoforms of AD-positively correlated modules (e.g., L1CAM_N671_ H3N4F1, NRCAM_223_H4N2, CNTN1_N338_H5N2, and OPCML_N70_H6N2) and AD-negatively correlated modules (CD200_N90_H6N6F1S2 and THY1_N42_H5N4F1 glycoforms) (Fig. 8A and table S9) supports a central role of cell adhesion glycoforms as key drivers of disease-associated glycoform network changes in AD. Our results (Fig. 7 and Fig. 8), together with increasing evidence for many of these cell adhesion proteins in controlling synapse organization and remodeling, synaptic function, trans-synaptic signaling, and/or neuron-glia communication (*44, 45*), indicate that dysregulated glycan modifications (e.g., up-regulated oligomannosylation and down-regulated sialylation) of cell adhesion proteins could be a previously unidentified pathogenic mechanism leading to synapse loss, synaptic dysfunction, and cognitive decline in AD.

Another prominent feature shared by all seven AD-related modules was their association with extracellular matrix (ECM) organization and matrisome function with an enrichment of glycoforms from matrisome proteins, including ECM structural constituents [e.g., laminins (LAMA2, LAMA5, and LAMC1), collagens (COL6A2 and COL6A3), heparan sulfate proteoglycan HSPG2/perlecan, fibromodulin (FMOD), nidogen-2 (NID2), biglycan (BGN), TINAGL1, and vitronectin (VTN)]; perineuronal net components [e.g., tenascins (TNC and TNR), neurocan (NCAN), aggrecan (ACAN), brevican (BCAN), hyaluronan and proteoglycan link protein 1 (HAPLN1), and versican (VCAN)], ECM receptors [e.g., integrins (ITGA1, ITGA2, ITGA6, ITGA7, ITGAM, ITGAV, ITGB1, and ITGB8), agrin (AGRN), BCAM, and tetraspanins (TSPAN3 and TSPAN8)], ECM-associated proteins [e.g., SPARCL1/hevin, clusterin (CLU), semaphorins (SEMA4D and SEMA7A), GPNMB/osteoactivin, and hemopexin (HPX)], and ECM regulators [ADAM10, lysyl hydroxylase-1 (PLOD1), A2M, alpha-1-antitrypsin (SERPINA1), and neuroserpin (SERPINI1)] (Fig. 8, D to F, and table S9). The importance of ECM glycoforms in driving AD-associated glycoform network changes is highlighted by their identification as top hub glycoforms of AD-positively correlated modules (e.g., LAMA2_N2028_H3N4F1, LAMA2_ N1901_H5N2, NCAN_N122_H6N2, HAPLN1_N56_H3N5F1, and LAMA5_N2501_H5N4F1S1) and AD-negatively correlated modules (LAMC1_N1107_H5N4F1, LAMC1_ N1223_H6N5F1S1, and TNR_N1261_H5N5F1S1) (Fig. 8A and table S9). Our findings (Fig. 7 and Fig. 8) reveal a previously unrecognized connection of aberrant mannosylation, fucosylation, and sialylation of matrisome proteins with altered ECM dynamics, matrisome dysfunction, and impaired ECM-cell communication in AD. Importantly, a number of the identified matrisome proteins (e.g., HSPG2, TNC, BGN, VCAN, ITGAV, ITGB1, ITGAM, CLU, SEMA7A, A2M, and SERPINA1) play critical roles in the control of inflammatory response (*46, 47*). Furthermore, AD-associated modules also contained glycoforms from other inflammation-related proteins [e.g., complement C3, complement factor H (CFH), HP, tissue factor F3, CD47, and JAM3]. Together, these results highlight a link between altered glycan modifications of these glycoproteins and neuroinflammation in AD.

Our analyses showed that AD-positively correlated GFM1, GFM2, and GFM4 modules were highly enriched with mannosylated and fucosylated glycoforms from neuronal and synaptic membrane proteins involved in neurotransmission and synaptic plasticity, including neurotransmitter receptors [glutamate receptors (GRM3/mGluR3, GRM5/mGluR5, GRIN1/NR1, GRIN2A/NR2A, GRIN2B/NR2B, GRIA1/GluR1, GRIA2/GluR2, GRIA3/GluR3, GRIK2/GluR6, and GRIK3/GluR7), GABA receptors (GABBR1, GABBR2, GABRA3, GABRB2, and GABRG2), and glycine receptor GPR158/mGlyR], synaptic vesicle proteins SYNPR and SYPL1, voltage-gated Ca^2+^ channel subunit CACNA2D1, neuronal K^+^/Cl^-^ cotransporter SLC12A5/KCC2, K^+^ channel modulatory subunit DPP6, glutamate receptor trafficking regulator OLFM1/noelin, and hippocampal excitatory synapse organizer IGSF8 with GRM3_N414_H3N4F1, GRIK2_ N378_H8N2, GRIN1_N276_H5N2, GABBR1_N482_H5N2, GRM5_N382_H3N5F1, and SYNPR_N33_ H3N5F1 as top hub glycoforms (Fig. 8, A, E, and F; and table S9), indicating an association of up-regulated mannosylation and fucosylation of the neurotransmitter receptors and synaptic proteins with synaptic dysfunction and cognitive decline in AD. By contrast, AD-negatively correlated GFM3 module was associated with the neurotransmission process with an enrichment of sialylated glycoforms from neuronal and synaptic proteins involved in ion transport, including Na^+^/K^+^ transporting ATPase subunits ATP1B1 and ATP1B2, voltage-gated Na^+^ channel subunit SCN2A/Nav1.2, glutamate ionotropic receptor subunit GRIA2/GluR2, synaptic vesicle proteins SV2A and SYPL1, K^+^ channel modulatory subunits DPP6 and DPP10, and choline transporter-like protein SLC44A2 with ATP1B1_N193_H7N6S3, ATP1B1_N193_H7N6F1S3, and SYPL1_N71_H6N4F1S1 as top hub glycoforms (Fig. 8, A, E, and F; and table S9), revealing an involvement of decreased sialylation of these proteins in altered brain cell ion homeostasis and neurotransmission in AD.

Another key finding from our analyses is a strong association of AD-positively correlated GFM1, GFM2, and GFM4 modules and AD-negatively correlated GFM3 module with signal transduction processes in the brain, as demonstrated by an enrichment of glycoforms from a variety of signaling receptors [e.g., insulin receptor (INSR), neurotrophic tyrosine kinase receptor NTRK2/TrkB, GDNF receptor GFRA2, EGF family receptors (EGFR and ERBB3), protein tyrosine phosphatase receptor PTPRZ1, purinergic receptor P2RX7, inositol 1,4,5-trisphosphate receptor ITPR1/IP3R1, neuronal pentraxin NPTX1, neuronal pentraxin receptor NPTXR, interleukin-6 receptor subunit IL6ST, V-set immunoregulatory receptor VSIR, adhesion G protein-coupled receptors (ADGRB1/BAI1, ADGRB3/BAI3, and ADGRL1/latrophilin-1), integrins (ITGA1, ITGA2, ITGA6, ITGA7, ITGAM, ITGAV, ITGB1, and ITGB8), and plexin family of semaphorin receptors (PLXNA1, PLXNA2, PLXNA4, PLXNB1, PLXNB2, and PLXNC1)], signaling receptor ligands (e.g., semaphorins SEMA4D and SEMA7A), and other signaling regulators (e.g., ADAM10, ADAM22, Nicastrin, ACE1, DKK3, MEGF8, NEGR1, LINGO1, and CD38) (Fig. 8E and table S9). These results, together with the identification of site-specific glycoforms from a number of signaling receptors and regulators as top hub glycoforms of AD-positively correlated modules (e.g., ADGRB3_N779_H5N2, NPTX1_N154_H3N5F1, ADAM22_N634_H3N5F1, NTRK2_N95_H6N2, and PLXNB2_N127_ H3N5F1) and AD-negatively correlated modules (e.g., LINGO1_N505_H3N6F1S2 and NEGR1_N294_H8N7F1), indicate that elevated mannosylation and fucosylation and reduced sialylation and N-glycan branching of the signaling receptors and regulators could be previously unknown pathogenic events leading to neuronal signaling dysregulation in AD.

We found that six of the seven AD-related glycoform modules (GFM1, GFM4, GFM11, GFM19, GFM3, and GFM14) were closely linked to lysosomal function with an enrichment of glycoforms from lysosomal proteins, including lysosomal lumenal hydrolases [e.g., cathepsins (CTSD and CTSL), tripeptidyl-peptidase TPP1, palmitoyl-protein thioesterase PPT1, lysosomal alpha-glucosidase GAA, lysosomal acid glucosylceramidase GBA, acid ceramidase ASAH1, prenyl-cysteine oxidase PCYOX1, beta-glucuronidase GUSB, iduronate 2-sulfatase IDS, steroid sulfatase STS, hexosaminidase subunit HEXA, sialic acid acetylesterase SIAE, and lysosomal carboxypeptidase C (PRCP)], lysosomal membrane-bound hydrolases (e.g., phospholipase PLD3 and lysosomal acid phosphatase ACP2), lysosomal transporters (e.g., vacuolar H^+^-transporting ATPase subunits ATP6AP1 and ATP6V0E2, and ATP-binding cassette transporter ABCA2), and other lysosomal membrane glycoproteins (e.g., LAMP1, LAMP2, and TMEM25) (Fig. 8D and table S9). Furthermore, site-specific glycoforms from several lysosomal glycoproteins (e.g., PLD3_N387_N1F1, PLD3_N387_H3N5F1, PCYOX1_N323_H5N2, TPP1_N313_H5N2, ABCA2_N14_H3N5F1, ABCA2_N1775_H9N2, and CTSL_N221_H6N2) were identified as top hub glycoforms of AD-related modules (Fig. 8A and table S9). These results, together with ample evidence for the critical roles of the identified lysosomal hydrolases, transporters, and other lysosomal glycoproteins in controlling intracellular degradation and brain homeostasis (*48, 49*) and the finding of PLD3 as a risk factor for AD (*50*), indicate that altered glycan modifications (e.g., increased paucimannosylation or oligomannosylation) of the lysosomal hydrolases and lysosomal membrane glycoproteins could be a previously unidentified pathogenic mechanism leading to lysosomal dysfunction, defective lysosomal acidification, impaired degradation, and neurodegeneration in AD.

Our analyses revealed that AD-positively correlated GFM1, GFM2, and GFM4 modules were also associated with the process of receptor-mediated endocytosis with an enrichment of mannosylated or fucosylated glycoforms from endocytic cargo receptors, including low density lipoprotein (LDL) receptor family members LRP1 and LRP1B, sortilin SORT1, sortilin-related receptor SORL1, and lysosomal sorting receptor M6PR/CD-MPR with LRP1_N1575_H5N2, LRP1_ N2620_H5N2, LRP1_N1511_H4N4F1, M6PR_N57_H7N2, and SORL1_N1894_H5N2 as top hub glycoforms (Fig. 8, E and F, and table S9). While fucosylated SORT1 glycoforms (SORT1_N98_H3N4F1, SORT1_N98_H4N5F1, and SORT1_N162_H3N5F1) were localized in AD-positively correlated GFM1 module, sialylated SORT1 glycoforms (SORT1_N98_ H4N5F1S1 and SORT1_N98_H4N6S1) were found in AD-negatively correlated GFM3 module linked to the process of receptor-mediated endocytosis (Fig. 8, E and F, and table S9). Furthermore, sialylated glycoforms from lysosomal sorting receptor SCARB2/LIMP2 (SCARB2_N224_H5N6F1S2 and SCARB2_N224_H6N6F1S2) were also localized in the GFM3 module (table S9). Our findings, together with the reported functions of these cargo receptors in controlling endocytic trafficking of APP, tau, and/or other cargos (*35, 51*) and GWAS identification of SORL1 and SORT1 as risk factors for AD (*52*), indicate that elevated mannosylation and fucosylation and reduced sialylation of the endocytic cargo receptors could be previously unrecognized pathogenic events contributing to endocytic trafficking dysregulation and AD neuropathology.

## Discussion

The present study is the first to use intact glycopeptide-based quantitative glycoproteomics coupled with network biology to provide a systems-level view of protein glycoforms and site-specific glycan modifications in human brain and their changes in AD. We report the largest human brain glycoproteome to date with 10,731 N-glycoforms from 1184 glycoproteins carrying 164 distinct glycan compositions across 2544 glycosylation sites. The acquisition of such extensive glycoproteomic information is mainly attributed to the strength of our optimized intact glyco-proteomics pipeline, including full extraction of proteins with strong detergent SDS (*25*), preparation of high-purity peptides by FASP (*26*), enrichment of intact glycopeptides using SAX-ERLIC, a more effective approach than lectin affinity enrichment methods for reducing the bias toward certain glycan moieties on glycopeptides (*27*), and SCE-HCD-based LC-MS/MS analysis (*28*). Our analyses reveal that oligomannosylation and fucosylation are two predominant types of glycan modification in human brain (Fig. 1, H and I, Fig. 3A, and table S4A), consistent with previous glycoproteomic and glycomic results (*24, 36*). Our study also shows that sialylation (including modifications by sialylated-only glycans and sialofucosylated glycans) is a common type of glycan modification in the brain, occurring at ∼26% of the total glycoforms (detection frequency) and ∼19% relative abundance (Fig. 1H, Fig. 3A, and table S4A). By contrast, a recent glycoproteomic study using multi-lectin and HILIC enrichment reported a low extent of sialylation at < 9% detection frequency in the brain (*24*), whereas a separate study using membrane fractions reported a predominance of sialofucosylation in the brain (*53*). The deep coverage of human brain glycoproteome achieved by our study provides fundamental information for understanding the roles of glycan modifications in brain health and disease.

Our deep intact glycoproteomic analysis extends our previous finding of altered N-glycosylation site occupancy in AD brain (*23*) and generates site-specific glyco-maps of N-glycans and glycoforms for human brain glycoproteins, including disease-relevant proteins such as GWAS-identified AD risk factors (e.g., CLU, SORL1, ABCA7, ACE, ADAM10, CNTNAP2, EPHA1, PLD3, and TM2D3) and other AD-related proteins (e.g., tau, nicastrin, and LRP1) in healthy and AD brains (Fig. 2, fig. S4, and table S2). The identified AD-associated changes in site-specific glycan modification patterns of these proteins provides compelling evidence supporting the proposed role of aberrant N-glycosylation in AD pathogenesis (*17, 18*). The site-specific glyco-maps of AD-relevant proteins obtained from this work provide an important resource for future studies to determine functional impacts of glycan modifications of these proteins in AD pathogenesis.

This study reveals that reduced sialylation is a prominent feature of AD brain glycoproteome (Fig. 1, H and I, Fig. 3A, and table S4A). Our finding of reduced sialylation in AD is in line with previously reported decrease in sialyltransferase activity in postmortem AD brain tissue samples (*54*). GWAS identification of sialic-acid-binding receptor CD33 (also called Siglec-3) as a risk factor for AD (*55*) underscores the importance of studying protein sialylation in brain dysfunction in AD. Our analysis shows that reduced overall sialylation in AD (Fig. 3A and table S4A) results mainly from disease-associated decreases in modification by 25 sialylated glycan compositions (Fig. 3, D and E). Moreover, we have identified 86 hyposialylated glycoforms from 39 hyposialylated proteins and 60 hypersialylated glycoforms from 44 hypersialylated proteins in AD (Fig. 5, A to C, and table S6, I and J) and uncovered multiple biological processes impacted by altered sialylation in AD, including cell adhesion, synaptic transmission, extracellular matrix function, cell surface receptor signaling, and lysosome function (Fig. 5, D to F, and table S7, I and J), providing mechanistic insights into the functional involvement of dysregulated protein sialylation in AD pathogenesis.

Another key finding from this work is the reduced N-glycan branching in AD brain. Our analyses have uncovered significant decreases in the overall levels of protein modifications by branched N-glycan with six or more HexNAc residues (Fig. 3B) and identified 21 glycan compositions containing ≥ 6 HexNAc residues as down-regulated glycan modifications in AD (Fig. 3, D and E, and table S4). Furthermore, out study reveals a list of glycoproteins and glycoforms with reduced N-glycan branching in AD (Fig. 4 and fig. S6). These results, together with the reported roles of N-glycan branching in controlling cell surface protein function, endocytosis, inflammation, and neurodegeneration (*56, 57*), indicate that decreased N-glycan branching is a previously unknown pathogenic mechanism contributing to endocytic trafficking dysregulation, neuroinflammation, and neurodegeneration in AD.

The observed predominant presence of high-mannose glycans on human brain glycoproteins (Fig. 1, H and I, and Fig. 3A) is consistent with prior studies showing that unlike most tissues, brain contains high abundance of high-mannose glycans, which are concentrated at the synapse (*58, 59*). Our study reveals that, while the overall levels of oligomannosylation by high-mannose glycans are unaltered in AD (Fig. 1, H and I, and Fig. 3A), H5N2 (Man5 glycan) modification is increased and H9N2 (Man9 glycan) modification is decreased in AD (Fig. 3, C to E). Furthermore, we have identified 187 glycoforms and 107 glycoproteins with hyperoligomannosylation and 36 glycoforms and 29 glycoproteins with hypo-oligomannosylation in AD (Fig. 5, A to C, and table S6, C and D). Our results indicate that disease-associated hyperoligomannosylation and hypo-oligomannosylation both affect plasma membrane and extracellular proteins involved in cell adhesion, transmembrane signaling, and inflammation in AD (Fig. 5, D to F, and table S7, C and D). However, only hyperoligomannosylation, but not hypo-oligomannosylation, is significantly associated with proteins localized to the synapse, ER, and lysosome to impact synaptic transmission, ER function, and lysosomal processes (Fig. 5, D to F).

This study shows that protein paucimannosylation is a significant feature of human brain glyco-proteome, occurring at ∼4% of the total glycoforms and ∼9% of the total glycosites (Fig. 1, H and I). Paucimannosylation was initially thought to be an invertebrate- and plant-specific type of glycosylation (*60*), but accumulating evidence indicates that paucimannosylation also occurs in vertebrates, particularly in the mammalian brain (*61*). Recent studies have shown that protein paucimannosylation is mediated by an N-acetyl-*β*-hexosaminidase-dependent truncation pathway (*62*) and paucimannose glycans on glycoproteins can be further truncated to form chitobiose-core-type glycans (*61*). Our finding of increased modifications by paucimannose glycans (H1N2, H2N2, and H3N2) and chitobiose-core-type glycans (N1 and N2) in AD brain (Fig. 3E and table S4C) indicates an involvement of elevated N-glycan truncation in AD. Corroborating with our finding, a glycomic study reported an association of AD neuropathology with a shift to smaller N-glycans (*63*). Our work reveals a list of glycoproteins and glycoforms with altered paucimannosylation in AD brain (Fig. 5, A to C, and table S6, E and F) and uncovers several hyperpaucimannosylation-affected processes in AD, including neuroinflammation, synaptic dysfunction, cell adhesion alteration, Golgi dysfunction, and cell signaling dysregulation (Fig. 5, D to F, and table S7E).

Our finding of a predominance of protein fucosylation in human brain (Fig. 1, H and I, and Fig. 3A) is in line with the reported high abundance of fucosylated N-glycans in the brain with crucial roles in regulation of synaptic plasticity and cognitive processes (*59, 64*). Our analyses reveal an increase in modification by fucosylated-only glycans and a decrease in modification by sialofucosylated glycans, resulting in no significant change in overall fucosylation in AD brain (Fig. 1, H and I, fig. S3, Fig. 3A, and table S4A). Furthermore, we have identified 188 glycoforms and 83 glycoproteins with hyperfucosylation and 83 glycoforms and 44 glycoproteins with hypofucosylation in AD (Fig. 5, A to C, and table S6, G and H). The elucidation of multiple biological processes and pathways impacted by disease-associated hyperfucosylation and hypofucosylation (Fig. 5, D to F, and table S7, G and H) provides evidence linking altered fucosylation to AD pathophysiology.

This work establishes glycan modification co-regulation network analysis as a new approach to analyze complex datasets of intact glycoproteomics. Unlike differential analysis which determines abundance changes of individual glycan modifications independently, co-regulation network analysis relates glycan modifications to one another using pairwise correlation relationships between glycan modification profiles to illuminate higher-order organization and define modules of co-regulated glycan modifications that are functionally related and/or coordinately regulated. Using this network analysis, we have uncovered an organization of human brain glycome (164 glycan compositions) into 8 modules of co-regulated glycan modifications (Fig. 6, A and B, and table S8). Importantly, we have identified 3 disease-associated glycan modules that are significantly correlated with AD clinicopathological phenotypes, including two negatively correlated modules (GNM1 and GNM4) enriched with sialylated glycans and highly branched glycans as well as one positively correlated module (GNM2) enriched with oligomannose, paucimannose, chitobiose-core glycans, bisecting GlcNAc-containing glycans, and undecorated complex/hybrid glycans with relatively simple, neutral structures (Fig. 6 and table S8). Our glycan network analysis reveals previously unknown co-regulation relationships among different glycan modifications in human brain (Fig. 6D and table S8) and highlights an involvement of glycan-specific elevations in oligomannosylation, undecorated complex/hybrid modification, bisection, paucimannosylation, and chitobiose-core modification and reductions in sialylation and N-glycan branching in AD pathophysiology. These results illustrate the value of glycan co-regulation network analysis for generating systems-level insights into glycan modifications in health and disease.

Another new approach established by this work is the intact glycoproteomics data-driven glycoform co-regulation network analysis, which reveals an organization of human brain glycoproteome into 21 modules of co-regulated protein glycoforms (Fig. 7 and table S9). The human brain glycoform network shows little overlap with our previously reported proteomic (*30*) and deglycoproteomic (*23*) networks, indicating that glycoform co-regulation, protein co-expression, and N-glycosylation site occupancy are controlled by distinct mechanisms. Our study has identified 7 disease-associated, glycoform modules with significant correlations to AD phenotypes, including 5 positively correlated modules (GFM1, GFM2, GFM4, GFM11, and GFM19) and 2 negatively correlated modules (GFM3 and GFM14) (Fig. 7 and fig. S8). Our analyses have generated a molecular blueprint of human brain glycoforms and site-specific glycan modification networks in AD brain and uncovered a number of dysregulated glycan modification-impacted processes and pathways, including alterations in cell adhesion and synapse organization, matrisome dysfunction, neuroinflammation, synaptic dysfunction, impaired brain cell ion homeostasis, cell signaling dysregulation, ER and Golgi dysfunction, endocytic trafficking dysregulation, and lysosomal dysfunction (Fig. 7, Fig. 8, table S9, and table S10). The identified disease-associated networks, pathways, and hub glycoforms provide attractive targets for AD biomarker and therapeutic development.

We note that current glycoproteomics methods can only provide glycan compositions as the site-specific glycan information, which limits insight into glycan structural diversity as a single glycan composition may represent multiple glycan structures (*65*). In addition, quantitative glyco-proteomics is limited by the occurrence of missing data, such as missing glycopeptide identifications or abundance values (*66*). Although our quantitative analyses have identified 677 altered glycoforms in AD (Fig. 5, A and B, and table S6, A and B), some important glycoform changes may have been missed as a result of excluding intact glycopeptides with missing values in > 50% of the samples in our analyses. Furthermore, the sample size used in this study only allowed detection of disease-associated glycosylation changes with a large effect size, and more subtle changes may have been missed (*67*). Despite these limitations, we anticipate that the methodologies and findings presented here will facilitate further studies of disease-associated glycoforms and site-specific glycan changes in AD.

In summary, our work has established the integration of intact glycopeptide-based quantitative glycoproteomics with network biology as a new paradigm for analyzing proteome-wide changes in site-specific glycan modifications in AD. This new approach can be broadly applied to study glycan modification alterations in other human diseases or in different stress conditions. Our comprehensive and quantitative glycoproteomics data of healthy human brains and AD patients have revealed previously unknown disease signatures of altered glycoforms and site-specific glycan modifications in AD and will serve as a valuable resource to facilitate future investigations into the roles of glycan modifications in normal brain physiology and AD pathophysiology. The identified glycan modification aberrations and brain glycoform and glycan network alterations provide new insights into AD pathogenesis and pave the way forward for developing glycosylation-based therapies and biomarkers to combat this devastating disease.

## Materials and Methods

### Human brain sample preparation

Postmortem brain tissues were acquired with informed consent from the donors or their family and provided by Emory Center for Neurodegenerative Disease Brain Bank. All experiments were conducted in accordance with the U.S. National Institutes of Health (NIH) guidelines for research involving human tissues and with the ethical standards and principles of the Declaration of Helsinki. For intact glycoproteomics analysis, we used the same sets of dorsolateral prefrontal cortex samples from AD and age-matched control cases as described in our previous proteomics and deglycoproteomics studies (*23, 30*). The case information and neuropathological data, including age, sex, disease status, disease duration, Aβ pathology CERAD score, neurofibrillary tangle Braak stage, ApoE genotype, and postmortem interval (PMI), can be found in table S1. Power analysis showed that the sample size used in this study (n = 16 AD and control cases) has > 80% power at a two-sided Type I error rate of 5% to detect an effect size of at least 1.6. For glycoproteomics sample preparation, human brain tissue (25 mg per AD or control case) was homogenized in 150 µl of lysis buffer containing 4% SDS, 100 mM DTT, and 100 mM Tris–HCl (pH 7.6) at 95°C, and proteins were extracted as we described previously (*25, 30*). Protein extracts were processed by following the filter-aided sample preparation (FASP) protocol with the Microcon 30-kDa centrifugal filter device (MRCF0R030, Merck) for detergent removal, cysteine alkylation by iodoacetamide, trypsin digestion, and peptide purification (*25, 26*). Peptide concentrations were determined by UV spectrometry as described (*30*), and purified peptides were dried under vacuum.

### Enrichment of intact glycopeptides by SAX-ERLIC

Purified peptides from each brain sample (200 µg of peptides per sample) were reconstituted in 60 µL of 50% acetonitrile (ACN) and 0.1% trifluoroacetic acid (TFA) in H_2_O. Intact glycopeptides were enriched by using electrostatic repulsion-hydrophilic interaction chromatography (ERLIC) with strong anion exchange (SAX) solid-phase extraction (SAX-ERLIC) as described (*27*) with minor modification. Briefly, SOLA-AX Solid Phase Extraction (SPE) cartridges (10 mg sorbent per cartridge, Thermo Fisher Scientific) were equilibrated three times with 1 mL of ACN, three times with 100 mM triethylammonium acetate (pH 7.0), and three times with loading buffer (95% ACN and 1% TFA in H_2_O). Peptides were diluted in 1 mL loading buffer and loaded onto the SPE cartridge, and the flow-through was collected and loaded again onto the cartridge. After washing the cartridge with 1 mL of loading buffer three times, the bound glycopeptides were eluted in two 600 µL aliquots of elution buffer (50% ACN and 0.1% TFA). The eluted glycopeptides were dried under vacuum and stored at -20 °C for further analysis.

### Intact glycopeptide-based mass spectrometric analysis

LC-MS/MS glycoproteomic analysis of intact glycopeptides was performed using a Q-Exactive HF-X Orbitrap mass spectrometer equipped with a nano-spray source and a nano-LC UltiMate 3000 high-performance liquid chromatography system (Thermo Fisher Scientific). Intact glycopeptides were separated by online reversed phase-HPLC fractionation on an in-house packed column (30 cm x 100 μm; packed with 3 μm Reprosil-Pur C18-AQ beads from Dr Maisch, Ammerbuch-Entringen, Germany), using a linear gradient from 3% to 40% solvent B over 120 min at a flow rate of 350 nL/min (mobile phase A, 2% ACN, 98% H_2_O, 0.1% FA; mobile phase B, 80% ACN, 20% H_2_O, 0.1% FA). The mass spectrometer was operated in the data-dependent acquisition (DDA) mode. Full MS scans were acquired from m/z 800 to 2,000 at a resolution of 60,000 at m/z 200 with an automatic gain control target of 1×10^6^ ions and maximum ion accumulation time of 50 ms. The 15 most abundant ions were selected from each full MS scan and fragmented with higher-energy collisional dissociation (HCD) using a stepped collision energy (SCE) setting of 18% / 28% / 38%. SCE-HCD-MS/MS spectra were acquired at a resolution of 15,000 at m/z 200 with an automatic gain control target of 20,000 ions and maximum ion accumulation time of 60 ms.

### Database search and intact N-glycopeptide identification

MS raw data were processed by searching against the UniProt human protein database (2016_02 Release, 20,198 reviewed entries) and a Byonic built-in glycan database of 182 human N-glycans using Byonic software version 3.11.3 (Protein Metrics) (*29, 68*). Searches were conducted with the fragmentation type set to HCD and mass tolerance setting of 20 ppm for both precursor and fragment ions. Trypsin was selected as the proteolytic enzyme with a maximum of two missed cleavages allowed. Carbamidomethylation of cysteine (+57.021464 Da) was set as a fixed modification, and the variable modifications included: N-glycosylation of asparagine residue localized within the sequon N-X-S|T (X ≠ P), oxidation of methionine (+15.994915 Da), and deamidation of asparagine and glutamine (+0.984016 Da). N-glycosylation and deamidation were set as “common” modification, and oxidation was set as “rare” modification. A total of two common modifications and one rare modification were allowed per identified peptide. The search output was filtered to <1% false discovery rate (FDR) using the FDR 2D score, a Byonic-computed FDR value based on a two-dimensional target-decoy strategy that estimates and controls peptide-spectrum match (PSM)- and protein-level FDRs simultaneously (*29, 69*). The filtered data were further assessed using the posterior error probability (PEP) 2D score, a comprehensive measure of error probability by taking into account ten different Byonic parameters, including the Byonic score, delta, precursor mass error, and digestion specificity (*29, 68*). Only the intact N-glycopeptides with FDR 2D score < 0.01 and PEP 2D score < 0.05 were included in data analysis.

### Quantitative analysis of intact glycopeptides and N-glycan modifications

Quantification of intact N-glycopeptides was performed by using Byologic software version 3.11.3 (Protein Metrics), which uses inputs from both MS1 raw data and Byonic search results to determine glycopeptide intensities (*68*). The Byologic quantification results were filtered to FDR 2D score < 0.01 and PEP 2D score < 0.05. For quantitative analysis of N-glycan modifications, intact glycopeptides were grouped by their glycan composition or glycan type, and the relative abundance of glycan modification for each glycan category was determined by the summed intensities of glycopeptides for the glycan category normalized by total intensities of glycopeptides for all N-glycan modifications in each sample. Significance of difference in relative abundances of glycan modification for each glycan category between AD and control brains was assessed using unpaired two-tailed Student’s *t* test with *P* < 0.05 as the confidence threshold. Unsupervised hierarchical clustering based on the relative abundances of glycan modifications in each samples was performed as we described previously (*30*).

### Differential analysis of glycoform abundance

The abundance of a site-specific glycoform was determined by summing the intensities of intact glycopeptides with the same glycan composition at the same glycosylation site within the same protein. Glycoform abundance values were converted to log_2_ scale and normalized by applying median-centered normalization. Differential abundance analysis was performed using the normalized abundances of glycoforms with valid abundance values in ≥ 50% of samples per group by employing an empirical Bayes approach with the *limma* R package (*70, 71*). Glycoforms with altered abundance in AD were identified by using *limma* moderated *t* test with the thresholds of ±1.3-fold change over the control (i.e., AD/control ratio > 1.3 or < 0.77) and *P* < 0.05. The *q* value was used to correct for multiple hypothesis testing and estimate the FDRs as described (*72*). Unsupervised hierarchical clustering based on the normalized glycoform abundances in each samples was performed as described (*30*), and the clustered cases and glycoforms were visualized along with the corresponding N-glycan types using the Heatmapper software (*73*). In addition to the quantitative analysis for identification of glycoforms with altered N-glycan modification levels, glycoforms with a gain of a site-specific N-glycan modification in AD were identified as the glycoforms detected in ≥ 50% of AD cases but not in any of the control cases, and glycoforms with a complete loss of a site-specific N-glycan modification in AD were identified as the glycoforms detected in ≥ 50% of control cases but not in any of the AD cases.

### Glycan modification co-regulation network analysis

Data-driven glycan modification co-regulation network analysis was performed by using WGCNA algorithm (*39*) to construct a human brain glycan network from the glycan modification abundance profiles of 164 glycan compositions. We used WGCNA package in R (*39*) to calculate a correlation matrix for all pairwise correlations of glycan modification abundances across all AD and control samples and then transformed it into a weighted adjacency matrix with a soft threshold power of 6 according to the scale-free topology criterion (*39*). The weighted adjacency matrix was used to generate a topological overlap (TO) matrix, and hierarchical clustering of glycans was performed using 1 − TO as the distance measure. Modules of co-regulated glycan modifications were identified using dynamic tree-cutting with the following parameters: minimal module size = 5, deepSplit = 4, merge cut height = 0.07, and a reassignment threshold of *P* < 0.05. Glycan co-regulation module networks were graphically depicted by using the Cytoscape v.3.5.0 software (*74*).

### Glycoform co-regulation network analysis

The intact glycoproteomics dataset was filtered for N-glycoforms with valid abundance values in ≥ 50% of AD or control samples, and the missing abundance values were imputed using the *k*-nearest neighbors imputation function in the DAPAR software (*75*). Glycoform co-regulation network analysis was performed using the WGCNA package in R (*39*) to generate a correlation matrix for all pairwise correlations of glycoform abundances across all samples followed by transformation into a weighted adjacency matrix using a soft threshold power of 10. Glycoforms were hierarchically clustered using the TO-based dissimilarity, and modules of co-regulated glycoforms were identified by dynamic tree-cutting with parameters set as minimal module size = 30, deepSplit = 4, merge cut height = 0.07, and a reassignment threshold of *P* < 0.05. Glycoform co-regulation module networks were graphically depicted by using the igraph software package (*76*).

### Analysis of inter-modular relationships and module-trait associations

For each module in the glycan or glycoform co-regulation network, we computed a module eigenglycan or eigenglycoform as the first principal component of the co-regulated glycans or glycoforms in the module according to the definition of eigennode (*39*). Pearson correlation between each glycan or glycoform and the module eigenglycan or eigenglycoform was used to determine module membership *kME*, a measure of intramodular connectivity (*42, 77*). Hub glycans or glycoforms for each module were identified as the glycans or glycoforms with *kME* > 0.8. Inter-modular relationships between glycan or glycoform co-regulation network modules were determined on the basis of pairwise correlations between module eigenglycans or eigenglycoforms as described (*77, 78*). Module-trait association analysis was performed as described (*30*) using biweight midcorrelations between each module eigenglycan or eigenglycoform and each clinical or neuropathological trait and the corresponding *P* values.

### Gene ontology and cell-type marker enrichment analyses

Gene ontology (GO) enrichment analysis of the generated datasets of differentially N-glycosylated proteins and glycoform network modules was performed using MetaCore bioinformatics software (version 6.37, build 69,500) as described (*30*), with the total list of all proteins (including glycoproteins) identified in our human brain samples as the background and FDR-corrected *P* <0.05 as the significance threshold. Enrichment analysis for markers of major brain cell types (astrocyte, microglia, neuron, and oligodendrocyte) in glycoform network modules was performed as we described previously (*23*). Significance was determined using one-sided Fisher’s exact test followed by correction for multiple comparisons using the Benjamini-Hochberg FDR adjustment method (*79*).

### Statistical Analyses

The sample size used in this study was determined based on power analysis and our prior studies (*23, 30*). Significance of difference in group means between AD and control cases was assessed by unpaired two-tailed Student’s *t* test, *limma* moderated *t* test, or Kruskal–Wallis test as indicated. For data visualization, heatmaps with Z-score values of normalized abundances were generated using the Heatmapper tool (*73*). Glycan or glycoform co-regulation network analysis was performed using the WGCNA algorithm (*39*) to define network modules of co-regulated glycans or glycoforms. Module-trait association was assessed by biweight midcorrelation between each module eigenglycan or eigenglycoform and each trait with *P* < 0.05 as the confidence threshold. Enrichment analyses were performed with one-sided Fisher’s exact test to calculate *P* values. Correction for multiple comparisons was performed using the *q* value (*72*) or Benjamini-Hochberg FDR adjustment method (*79*) as indicated.

## Acknowledgments

We thank Dr. Marla Gearing for providing postmortem brain tissue samples with demographic and neuropathological information. We are grateful to the Emory Center for Neurodegenerative Disease Brain Bank donors and their families for their invaluable contribution to this study. S.P. is grateful to the endowment of the Rochelle and Max Levit Chair in the Neurosciences.

## Author contributions

L.L. and Q.Z. conceived and designed the experiments. Q.Z. and C.M. performed sample preparation and glycoproteomics analyses. Q.Z. performed glycopeptide quantification and data analyses. Q.Z. and L.L. performed bioinformatics and network analyses. L.L., L.S.C., and S.P. acquired funding and provided supervision. L.L., Q.Z., and L.S.C. drafted the manuscript. All authors edited, reviewed, and approved the manuscript.

## Data and materials availability

The mass spectrometry glycoproteomics data have been deposited to the ProteomeXchange Consortium via the PRIDE (*80*) partner repository with the dataset identifier PXD045001. All data needed to evaluate the conclusions in the paper are present in the paper and/or the Supplementary Materials.

