## Supplementary Materials for "Human brain glycoform co-regulation network and glycan modification alterations in Alzheimer’s disease"

Qi Zhang *et al*

**The PDF file includes:**

Figs. S1 to S8
Table S1

**Other Supplementary Material for this manuscript includes the following:**

Table S2 to S10 (Microsoft Excel format)

**SUPPLEMENTARY FIGURES**

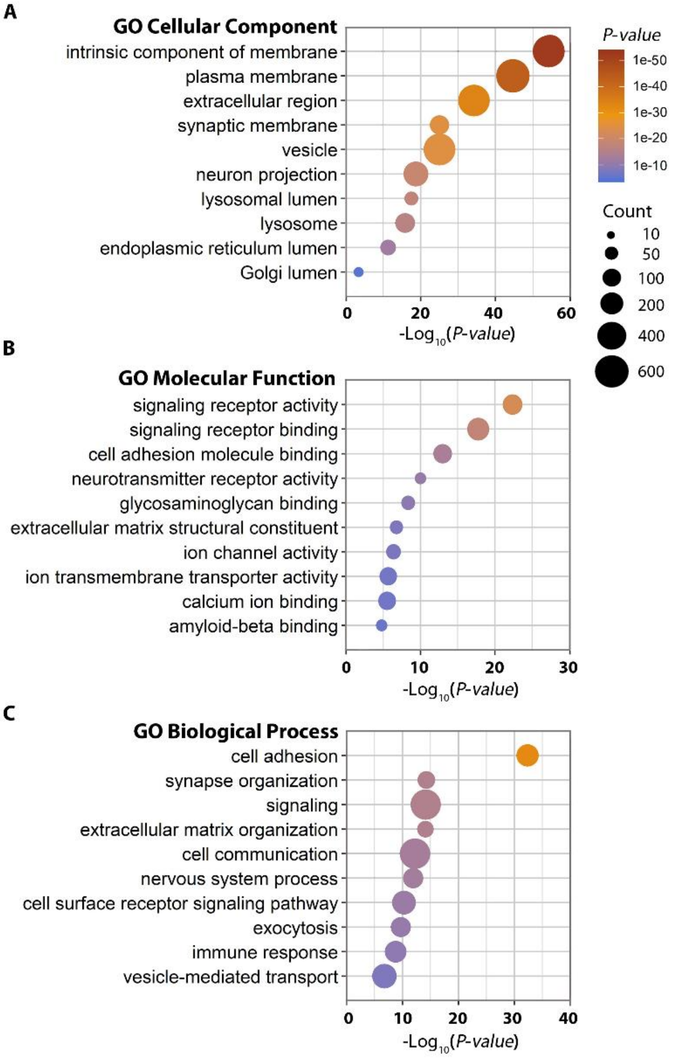


**Fig. S1. Gene ontology enrichment analysis of human brain N-glycoproteins identified by intact glycoproteomics.** GO cellular component (**A**), molecular function (**B**), and biological process (**C**) enriched in the intact glycoproteome dataset are shown with Benjamini-Hochberg FDR-corrected *P* values. Count indicates the number of glycoproteins per GO term.

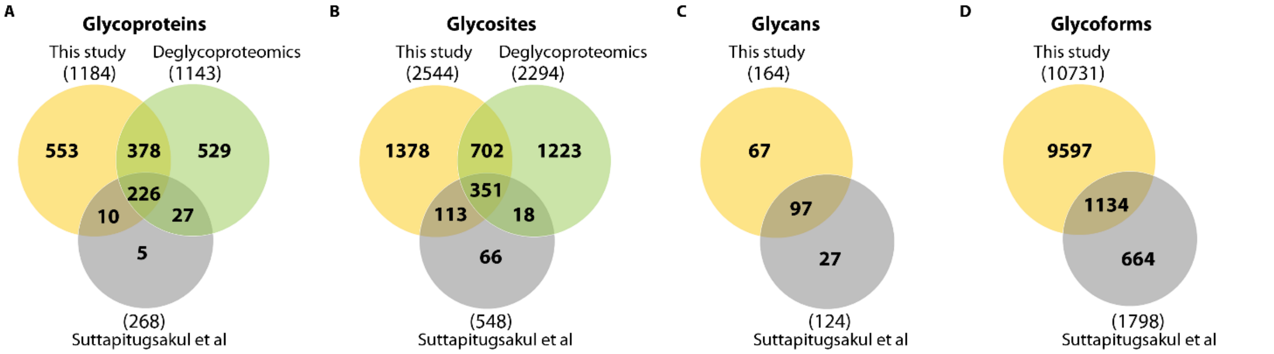


**Fig. S2. Comparison of human brain glycoproteomics dataset obtained in this study to Suttapitugsakul et al dataset and our deglycoproteomics dataset.** Venn diagrams showing the overlap of N-glycoproteins, N-glycosites, N-glycans, and N-glycoforms identified in this study with those from Suttapitugsakul et al (*Ref. 24*) and our deglycoproteomics study (*Ref. 23*).

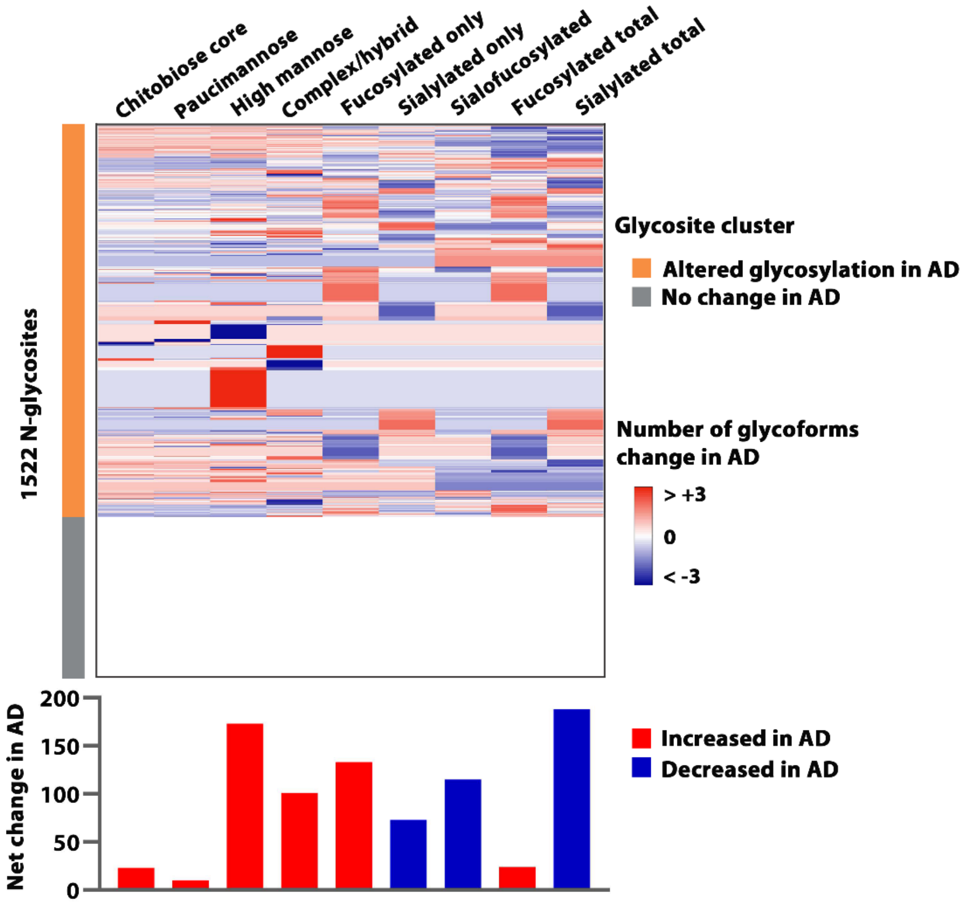


**Fig. S3. Disease-associated changes in glycosylation patterns of 1522 N-glycosites identified in both control and AD brains.** Heatmap (*top*) shows the unsupervised hierarchical clustering of 1522 N-glycosites based on site-specific changes in the number of glycoforms in AD versus the control for each of the indicated glycan types. The net change in the number of glycoforms for each glycan type across the 1522 glycosites in AD versus the control is shown at the *bottom*.

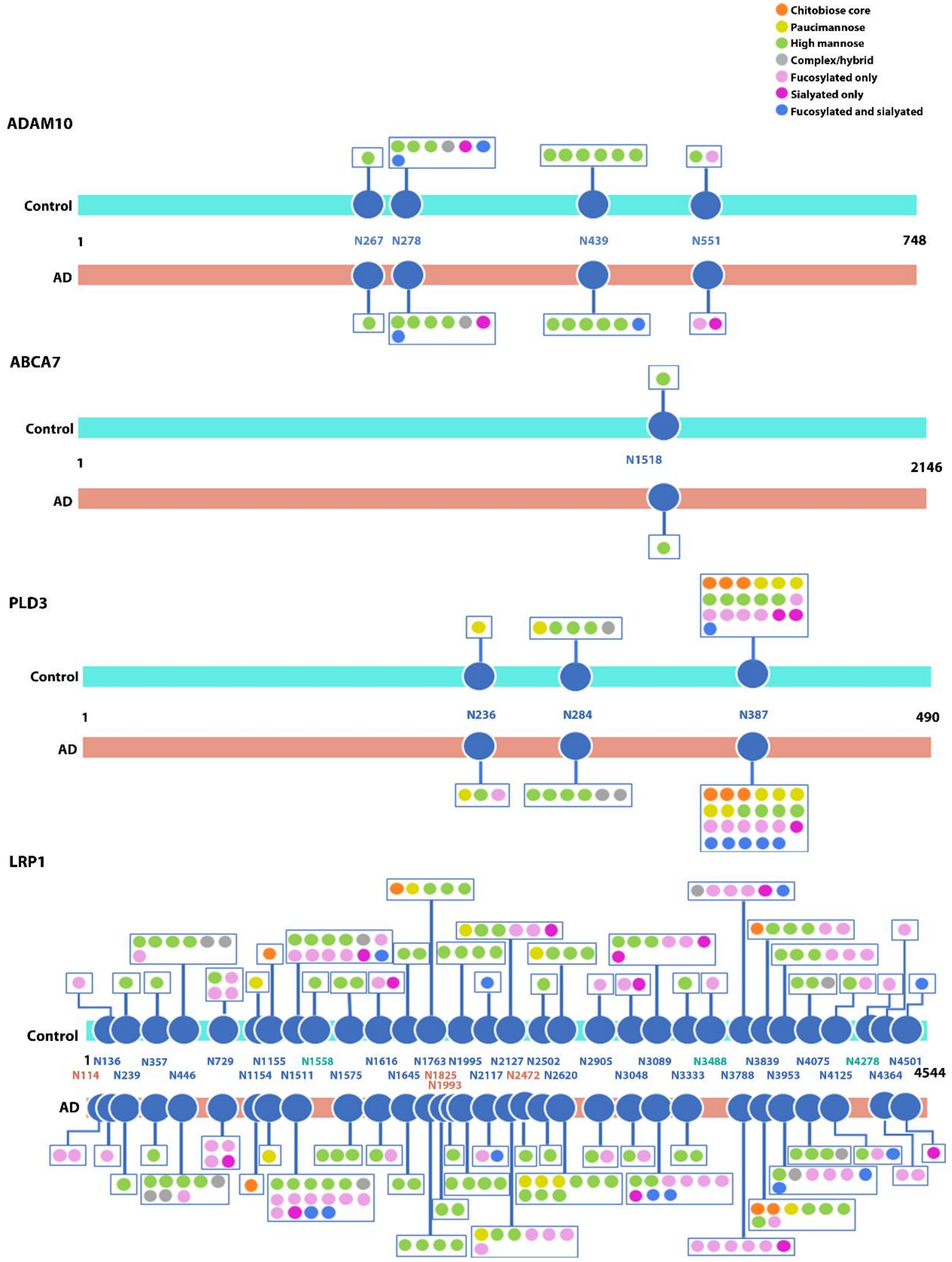


**Fig. S4. Site-specific glycosylation patterns of ADAM10, ABCA7, PLD3, and LRP1 in AD and control brains.** N-glycosites are indicated by the amino acid positions of glycosylated asparagine residues, with cyan- and pink-colored glycosites denoting the N-glycosites detected exclusively in control and AD brains, respectively.

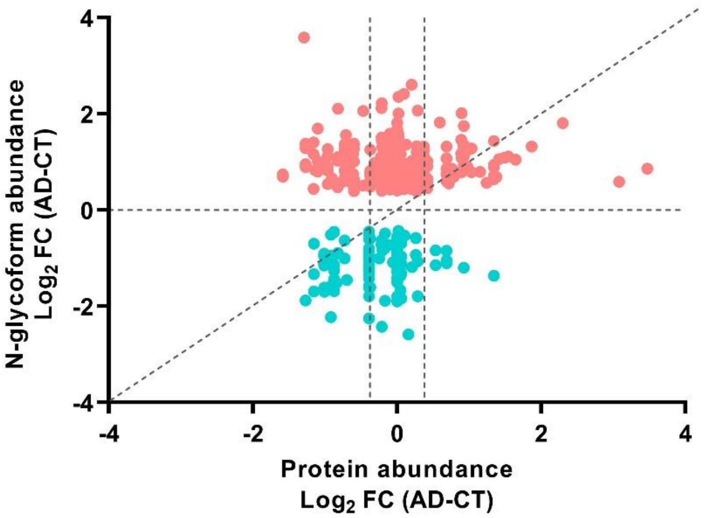


**Fig. S5. Comparison of glycoform abundance changes in AD with corresponding protein abundance changes.** Scatter plot showing that glycoform abundance changes of many AD-associated N-glycoforms are not due to altered protein abundance.

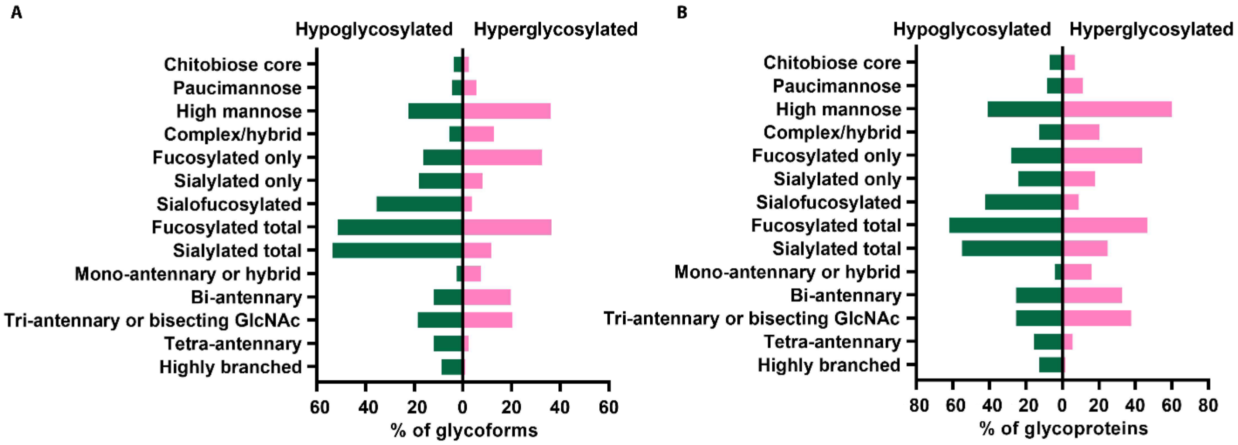


**Fig. S6. Distribution of glycoforms and glycoproteins by glycan type in AD hyperglycosylation and hypoglycosylation datasets.** Bar graphs showing the percentages of AD-associated hypoglycosylated or hyperglycosylated glycoforms (A) and glycoproteins (B) carrying the indicated glycan types.

**Glycan module-trait relationships**

| | Age | Sex | AD status | CERAD | Braak | ApoE | PMI |
|---|---|---|---|---|---|---|---|
| GNM8 | 0.23 (0.4) | 0.27 (0.3) | -0.22 (0.4) | -0.32 (0.2) | -0.15 (0.6) | -0.097 (0.7) | 0.25 (0.4) |
| GNM4 | -0.14 (0.6) | -0.14 (0.6) | -0.52 (0.04) | -0.57 (0.02) | -0.45 (0.08) | -0.25 (0.4) | 0.0057 (1) |
| GNM3 | 0.041 (0.9) | 0.16 (0.6) | -0.38 (0.1) | -0.4 (0.1) | -0.42 (0.1) | 0.31 (0.2) | 0.5 (0.056) |
| GNM1 | -0.0014 (1) | 0.23 (0.4) | -0.9 ($3x10^{-6}$) | -0.87 ($1x10^{-5}$) | -0.9 ($2x10^{-6}$) | -0.24 (0.4) | 0.31 (0.2) |
| GNM7 | 0.073 (0.8) | -0.0032 (1) | 0.13 (0.6) | 0.15 (0.6) | 0.042 (0.9) | -0.071 (0.8) | 0.28 (0.3) |
| GNM5 | -0.073 (0.8) | 0.35 (0.2) | 0.14 (0.6) | 0.2 (0.5) | 0.047 (0.9) | 0.26 (0.3) | 0.36 (0.2) |
| GNM2 | 0.042 (0.9) | -0.28 (0.3) | 0.92 ($3x10^{-7}$) | 0.91 ($1x10^{-6}$) | 0.85 ($4x10^{-5}$) | 0.29 (0.3) | -0.15 (0.6) |
| GNM6 | -0.36 (0.2) | -0.16 (0.6) | 0.24 (0.4) | 0.25 (0.4) | 0.23 (0.4) | -0.19 (0.5) | -0.021 (0.9) |

**Fig. S7. Module-trait relationships of glycan network modules to AD-related traits or sample characteristics.** Biweight midcorrelations between module eigenglycans and the indicated trait are shown with the correlation coefficients (bicor *r*-values) on the top and *P* values in the bracket below. Significant positive correlations ($r > 0.50$, $P < 0.05$) are highlighted in *Red*, and significant negative correlations ($r < -0.50$, $P < 0.05$) in *Green*. PMI, postmortem interval.

**Glycoform module-trait relationships**

| | Age | Sex | AD status | CERAD | Braak | ApoE | PMI |
|---|---|---|---|---|---|---|---|
| GFM3 | 0.071 (0.08) | 0.2 (0.5) | -0.92 ($4x10^{-7}$) | -0.88 ($7x10^{-6}$) | -0.91 ($1x10^{-6}$) | -0.25 (0.4) | 0.24 (0.4) |
| GFM10 | 0.48 (0.06) | -0.26 (0.3) | -0.37 (0.2) | -0.39 (0.1) | -0.47 (0.07) | 0.28 (0.3) | -0.035 (0.9) |
| GFM13 | 0.47 (0.1) | 0.11 (0.7) | -0.39 (0.1) | -0.34 (0.2) | -0.49 (0.06) | 0.088 (0.7) | 0.13 (0.6) |
| GFM17 | 0.43 (0.1) | 0.15 (0.6) | -0.12 (0.7) | -0.13 (0.6) | -0.21 (0.4) | 0.38 (0.1) | 0.36 (0.2) |
| GFM14 | 0.4 (0.1) | 0.46 (0.07) | -0.68 (0.004) | -0.66 (0.005) | -0.73 (0.001) | -0.19 (0.5) | 0.45 (0.08) |
| GFM20 | 0.41 (0.1) | 0.54 (0.03) | -0.028 (0.9) | -0.1 (0.7) | -0.073 (0.8) | 0.0055 (1) | 0.31 (0.2) |
| GFM9 | 0.14 (0.6) | 0.38 (0.1) | 0.24 (0.4) | 0.17 (0.5) | 0.29 (0.3) | 0.07 (0.8) | 0.15 (0.6) |
| GFM15 | -0.11 (0.7) | -0.093 (0.7) | 0.29 (0.3) | 0.26 (0.3) | 0.25 (0.4) | 0.21 (0.4) | -0.19 (0.5) |
| GFM16 | 0.39 (0.1) | 0.2 (0.5) | -0.18 (0.5) | -0.16 (0.6) | -0.2 (0.4) | 0.25 (0.4) | 0.29 (0.3) |
| GFM7 | 0.3 (0.3) | 0.17 (0.5) | -0.18 (0.5) | -0.11 (0.7) | -0.27 (0.3) | -0.058 (0.8) | 0.14 (0.6) |
| GFM5 | 0.058 (0.8) | 0.058 (0.8) | 0.17 (0.5) | 0.18 (0.5) | 0.1 (0.7) | -0.25 (0.3) | -0.13 (0.6) |
| GFM21 | 0.055 (0.8) | -0.13 (0.6) | 0.16 (0.5) | 0.23 (0.4) | 0.029 (0.9) | 0.0089 (1) | -0.38 (0.1) |
| GFM2 | -0.091 (0.7) | -0.27 (0.3) | 0.62 (0.01) | 0.68 (0.003) | 0.48 (0.06) | 0.43 (0.1) | -0.14 (0.6) |
| GFM8 | -0.027 (0.9) | 0.051 (0.9) | 0.42 (0.1) | 0.41 (0.1) | 0.3 (0.3) | 0.22 (0.4) | -0.093 (0.7) |
| GFM19 | -0.45 (0.08) | -0.17 (0.5) | 0.72 (0.002) | 0.72 (0.002) | 0.63 (0.006) | -0.035 (0.9) | -0.31 (0.2) |
| GFM4 | -0.34 (0.2) | -0.43 (0.09) | 0.7 (0.002) | 0.67 (0.004) | 0.66 (0.006) | -0.1 (0.7) | -0.45 (0.08) |
| GFM12 | 0.14 (0.6) | -0.078 (0.8) | -0.23 (0.4) | -0.34 (0.2) | -0.16 (0.5) | $8x10^{-4}$ (1) | 0.15 (0.6) |
| GFM18 | 0.4 (0.1) | -0.3 (0.3) | -0.12 (0.7) | -0.16 (0.6) | -0.19 (0.5) | 0.41 (0.1) | 0.24 (0.4) |
| GFM1 | -0.12 (0.7) | -0.33 (0.2) | 0.9 ($2x10^{-6}$) | 0.86 ($2x10^{-5}$) | 0.84 ($4x10^{-5}$) | 0.53 (0.03) | -0.16 (0.6) |
| GFM11 | 0.016 (1) | -0.45 (0.08) | 0.52 (0.04) | 0.43 (0.1) | 0.53 (0.04) | 0.56 (0.02) | 0.074 (0.8) |
| GFM6 | 0.34 (0.2) | -0.2 (0.5) | 0.27 (0.3) | 0.21 (0.4) | 0.22 (0.4) | 0.64 (0.008) | 0.22 (0.4) |

**Fig. S8. Module-trait relationships of protein glycoform network modules to AD-related traits or sample characteristics.** Biweight midcorrelations between module eigenglycoforms and the indicated trait are shown by bicor *r*-values on the top and *P* values in the bracket below. Significant positive ($r > 0.50$, $P < 0.05$) or negative ($r < -0.50$, $P < 0.05$) correlations are highlighted. PMI, postmortem interval.

### SUPPLEMENTARY TABLES

#### Table S1. Case information of human AD and control brain samples

| Case | Sex | Age at death (yr) | Age at onset (yr) | Disease duration (yr) | Braak stage | CERAD score | ApoE genotype | PMI (hr) |
|---|---|---|---|---|---|---|---|---|
| Control | | | | | | | | |
| CT1 | Male | 65 | | | 0 | Sparse | E3/3 | 6 |
| CT2 | Female | 75 | | | I | None | E3/3 | 6 |
| CT3 | Male | 61 | | | II | None | E3/4 | <12 |
| CT4 | Female | 74 | | | II | None | E3/3 | 7 |
| CT5 | Male | 59 | | | I | None | E2/3 | 6 |
| CT6 | Female | 78 | | | II | None | E3/3 | 11.5 |
| CT7 | Male | 94 | | | II | None | E3/3 | 5.5 |
| CT8 | Female | 61 | | | II | None | n.d. | 6 |
| Alzheimer's disease | | | | | | | | |
| AD1 | Male | 79 | 73 | 6 | V-VI | Frequent | E4/4 | 6 |
| AD2 | Female | 72 | 59 | 13 | VI | Frequent | E3/4 | 7 |
| AD3 | Male | 67 | 56 | 11 | VI | Frequent | E2/3 | 6.5 |
| AD4 | Male | 77 | 70 | 7 | VI | Frequent | E3/4 | 12 |
| AD5 | Male | 74 | 60 | 14 | VI | Frequent | E3/3 | 2.5 |
| AD6 | Male | 68 | 60 | 8 | VI | Frequent | E3/3 | 14.5 |
| AD7 | Female | 61 | 51 | 10 | VI | Frequent | E3/3 | 6 |
| AD8 | Male | 69 | 59 | 10 | VI | Frequent | E3/4 | 3.5 |

Files not embedded in the Word file:

Table S2 to S10 (Microsoft Excel format)

**Captions for tables S2 to S10**

Table S2 (Microsoft Excel format). Intact N-glycopeptides, N-glycoproteins, N-glycosites, N-glycans, and N-glycoforms identified in AD and control brains.

Table S3 (Microsoft Excel format). Gene ontology (GO) enrichment analysis of human brain glycoproteins identified by intact glycoproteomics.

Table S4 (Microsoft Excel format). Quantitative analysis of N-glycan modification levels by glycan type (A) or glycan composition (B) in AD and controls and identified glycan compositions with altered N-glycan modification levels in AD brain (C).

Table S5 (Microsoft Excel format). Differential glycoform abundance analysis (A) and identified glycoforms with altered site-specific N-glycan modification levels (B) or with a complete loss or gain of N-glycan modification (C) in AD.

Table S6 (Microsoft Excel format). Identified glycoforms and glycoproteins with hyperglycosylation (A), hypoglycosylation (B), hyperoligomannosylation (C), hypo-oligomannosylation (D), hyperpaucimannosylation (E), hypopaucimannosylation (F), hyperfucosylation (G), hypofucosylation (H), hypersialylation (I), or hyposialylation (J) in AD.

Table S7 (Microsoft Excel format). GO enrichment analysis of glycoproteins with hyperglycosylation (A), hypoglycosylation (B), hyperoligomannosylation (C), hypo-oligomannosylation (D), hyperpaucimannosylation (E), hypopaucimannosylation (F), hyperfucosylation (G), hypofucosylation (H), hypersialylation (I), or hyposialylation (J) in AD.

Table S8 (Microsoft Excel format). Glycan modification co-regulation network analysis.

Table S9 (Microsoft Excel format). Protein glycoform co-regulation network analysis.

Table S10 (Microsoft Excel format). GO enrichment analysis of glycoproteins in AD-associated glycoform modules.
